# Improved model building for cryo-EM maps using local attention and 3D rotary position embedding

**DOI:** 10.1101/2024.11.13.623164

**Authors:** Baoquan Su, Kun Huang, Zhenling Peng, Alexey Amunts, Jianyi Yang

**Affiliations:** MOE Frontiers Science Center for Nonlinear Expectations, State Key Laboratory of Cryptography and Digital Economy Security, Research Center for Mathematics and Interdisciplinary Sciences, Shandong University, Qingdao 266237, China; School of Life Science, Westlake University, Hangzhou 310030, China; University of Münster, Schlossplatz 8, 48143 Münster, Germany; Department of Structural Biochemistry, Max Planck Institute of Molecular Physiology, Otto-Hahn-Str. 11, 44227 Dortmund, Germany

**Author notes:** Corresponding authors: JY, AA, ZP.

## Abstract

Constructing atomic models from cryogenic electron microscopy (cryo-EM) density maps is essential for interpreting molecular mechanisms. In this study, we present CryFold, an approach for *de novo* model building for cryo-EM maps, leveraging recent advancements in AlphaFold2 (1) to improve the state-of-the-art method ModelAngelo (2). To accommodate the cryo-EM map information, CryFold replaces the global attention mechanism in AlphaFold2 with local attention, which is further enhanced by a novel 3D rotary position embedding. CryFold produces more complete models, reduces the resolution requirement, and accelerates the modeling. The application of CryFold to three new maps with unknown structure demonstrates its ability to accurately distinguish between paralog sequences in noisy regions, detect previously uncharacterized proteins with unknown functions, precisely compartmentalize the map to isolate non-protein components, and improve the modeling of conformational changes. A particular case includes a 104-protein complex that has been modeled within a few hours, and a minor conformational change of a single protein domain has been detected at the periphery when models from two different maps were compared. CryFold stands as an accurate method currently available for model building of proteins in cryo-EM structure determination. The source code and model parameters are available at https://github.com/SBQ-1999/CryFold.

## Introduction

With the advent of direct electron detectors, single-particle cryo-EM has become the primary method for resolving biomolecular structures (3-5). A critical step in cryo-EM structure determination is the construction of atomic models from 3D density maps. The pace of model deposition derived from cryo-EM maps in the Protein Data Bank (PDB) has increased by two orders of magnitude, from 67 in 2012 to 5788 in 2024 (6). The exponential growth has been driven by advances in automation of data collection and processing (7), accessible software and web platforms empowering broader adoption (8), and community efforts to further democratize and improve cryo-EM methodologies (9), including the use of artificial intelligence (AI).

Previously, model building relied heavily on manual efforts, involving fitting homology models generated by tools like I-TASSER (10) into the density maps using graphic software, such as *Coot (11)* or Isolde (12). The process of generating a 3D reconstruction involves averaging thousands of non-identical particle projections, resulting in cryo-EM maps that consist of regions with variable resolutions. Consequently, areas with higher flexibility or partial occupancy are more challenging to interpret with precision. For instance, it took around six months to develop modeling and refinement protocols to build a model for the cryo-EM map (EMD-2566, overall resolution 3.2 Å) of the large subunit from the yeast mitoribosome with 13 new proteins (4). This was the first fully refined cryo-EM model of an asymmetric object that also illustrated cryo-EM’s power in resolving structures of large complexes isolated from native sources, where not all protein sequences are known, marking a turning point in structural biology ^5^. Today, individual components of complexes are routinely identified directly from the density maps, revealing novel regulatory pathways. In our hands, it led to the description of biochemical connection between biogenesis and fatty acid and iron-sulfur synthesis (13), evolutionary mechanisms for gene transfer from organelles to the nucleus (14), identification of gene splitting (15) and even the discovery of a new type of translational frameshift (16).

The amino acid sequence can be inferred directly from the side-chain densities, allowing for searches of protein-sequence databases (17). Tools like *FindMySequence*, which uses main-chain information to fit predicted residue-type probabilities, can query sequence databases of user’s choice (18). However, due to the labor-intensive nature of the process, many deposited PDB structures contain errors, limiting their biological interpretations (19, 20). This is particularly true when reliable reference structures are unavailable or when dealing with density maps at resolutions lower than 4 Å. As a result, model building becomes a bottleneck, highlighting the necessity of automating this process.

Several methods have been developed to derive atomic structures from cryo-EM density maps. These can be categorized into two groups: traditional and utilizing deep learning. Traditional model-building methods include Pathwalking (21, 22), Buccaneer (23), EM-Fold (24), Rosetta (25), MAINMAST (26), Phenix (27). In Pathwalking and MAINMAST, constructing all-atom structures from density maps is treated as a minimization problem, solved using optimization algorithms. Rosetta and EM-Fold first deduce fragment structures from the density map and assemble them using Monte Carlo sampling.

Deep learning-based methods encompass DeepMainmast (28), EMBuild (29), CR-I-TASSER (30), DeepTracer (31), ModelAngelo (2), Cryo2Struct (32). These approaches typically start by applying a deep network to detect the backbone structure from the density map. They diverge in their subsequent strategies for constructing all-atom models. For instance, DeepMainmast and EMBuild fit predicted structure models to the backbone to create all-atom structures. CR-I-TASSER integrates backbone information into I-TASSER’s (10) classical assembly simulations to generate all-atom models. DeepTracer connects backbone atoms by solving the traveling salesman problem before adding side-chain atoms for the complete structure. ModelAngelo, regarded as state of the art, uses predicted amino acid probabilities, refines backbone atoms with a graph-based network and constructs the all-atom model through Invariant Point Attention, followed by Hidden Markov Model (HMM)-based sequence correction. Cryo2Struct employs HMM and Viterbi algorithms to align the input sequences with predicted ones to build the all-atom structure. By incorporating components of protein structure prediction framework, current methods have achieved more accurate results in interpreting density maps (33).

Recently, a breakthrough came from deep learning based structure predictors, such as AlphaFold2 (AF2) (1), RoseTTAFold (34) and trRosetta (35), which also contributed to the awarding of the 2024 Nobel Prize in Chemistry. We reasoned that accuracy and the resolution limit in protein model building, especially for the state-of-the-art method ModelAngelo, could be further extended by taking advantage of the recent advances. We thus developed CryFold. Tests on cryo-EM maps of multi protein complexes with unknown components demonstrate the robustness of CryFold for automated model building with high speed and accuracy.

### Overview of the CryFold approach

CryFold takes as input a cryo-EM density map and the amino acid sequences of the target proteins, following a two-step process for building atomic structures (Fig. 1). In the first step, a 3D convolution-based network, specifically a U-Net (36), is employed to predict the positions of Cα atoms from the density map. Due to GPU memory limitations, the map is cropped into smaller patches (64×64×64), which are later combined to reconstruct the original size. The second step generates all-atom structures using novel transformer network adapted from AF2 (1). The all-atom positions are post-processed using a method similar to that used in ModelAngelo (2). CryFold incorporates a recycling mechanism, allowing for three iterative refinements of predictions like AF2 (1), and we introduced key modifications in the encoder and decoder to fully exploit the structural information present in the density map. The encoder, which we call Cryformer, transforms the density map into the node representation and the edge representation. The decoder, namely Structure Module, generates all-atom positions from these improved representations.

**Fig. 1.**
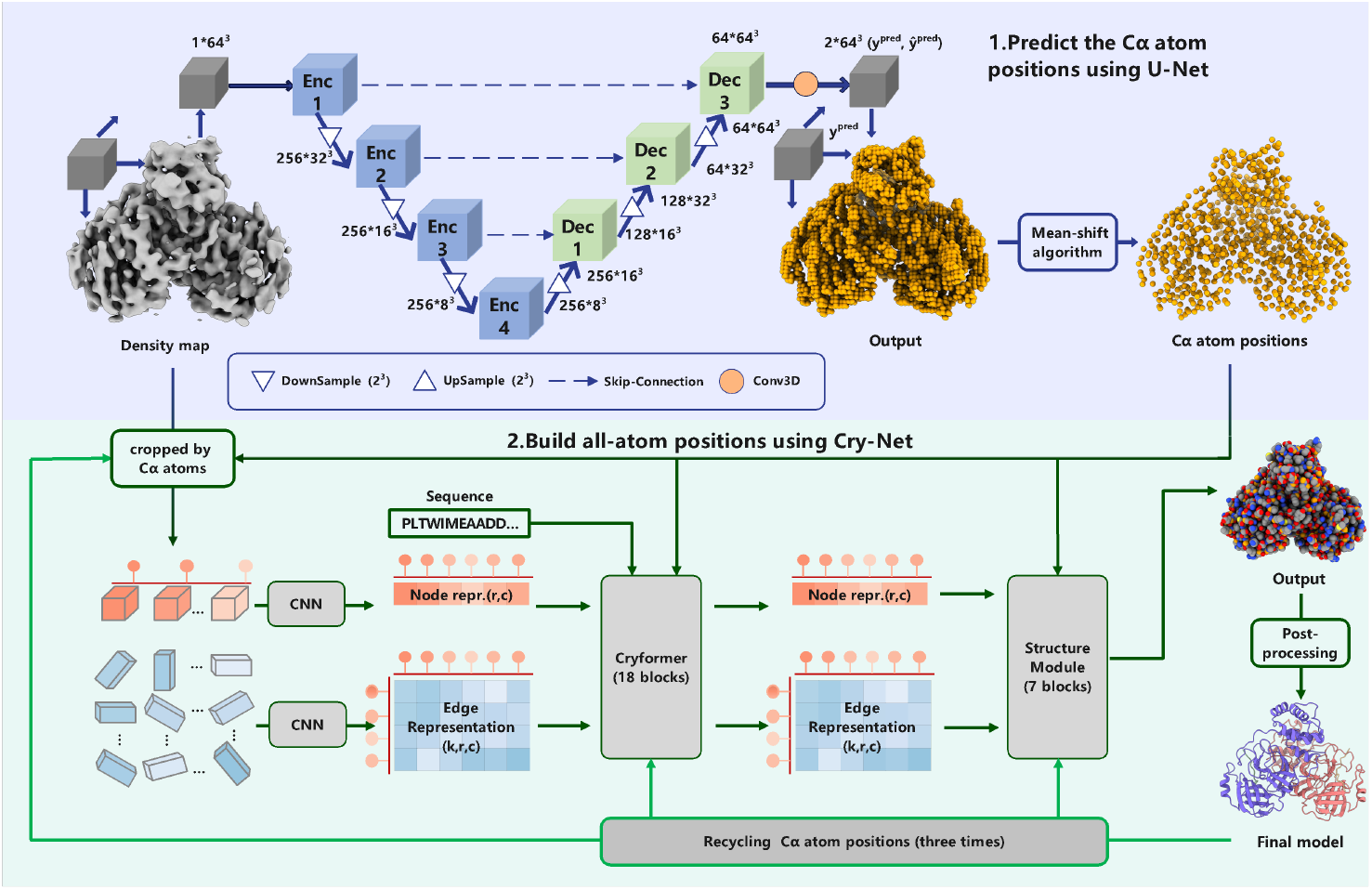
Overall architecture of CryFold. Two major steps are employed to build all-atom structure from a cryo-EM density map and the amino acid sequences of the target proteins. In the first step (with purple background), the input density map is fed into a 3D convolution-based network U-Net to predict the Cα atom positions. The map is first cropped continuously into boxes of size 64×64×64, which are then processed by U-Net and combined at the end. The final U-Net output *y*^*pred*^ is of the same size to the input map and the value in each voxel is the probability of containing a Cα atom, which is refined by the mean-shift algorithm. In the second step (with green background), all-atom positions are predicted using a local attention network (Cry-Net) from the amino acid sequence and the predicted Cα atom. The initial node and edge representations are first fetched from density map guided by the predicted Cα atom positions. They are then fed into the Cryformer together with the sequence embedding from ESM-2. The updated representations are then decoded into all-atom structure using the Structure Module, followed by a post-processing step. The parameters *r, k* and *c* are the numbers of predicted Cα atoms, neighbors considered and channels, respectively.

Two key differences distinguish the CryFold network (denoted by Cry-Net) from the AF2 network: an enhanced transformer and the local attention. Cry-Net features an enhanced transformer that incorporates 3D rotary position embedding (*3D*-*RoPE*). Originally, RoPE was used to process 1D sequences (37). Here we extend it to a node position encoding in 3D space (see Methods and Supplementary S2.4). 3D-RoPE effectively encodes the positional information of each node into all attention calculations, making the attention score to decay with increasing distance between nodes. This design takes advantage of the spatial constraints inherent in the density map. The local attention in Cry-Net is realized through the improved edge representation (see Methods). Instead of an all-against-all approach, Cry-Net employs an all-against-*k* strategy, where *k* represents the number of spatial neighbors for each node. This adjustment once again capitalizes on the spatial restraints provided by the density map, which are usually unavailable in traditional structure prediction. These features enable efficient training of a deeper network.

The post-processing procedure, adapted from ModelAngelo, is outlined as follows. The initial all-atom structure is obtained from Cry-Net, and any segments with fewer than four residues are trimmed (referred to as *model_net*). The amino acid type for each residue in model_net is assigned as the class with the highest probability output from Cry-Net. HMM profile is derived from the predicted probability distribution by Cry-Net and aligned with the input amino acid sequence. The amino acid types of residues in the Cry-Net structure (prior to trimming) are then corrected according to this alignment, producing another structure denoted as *model_fix*. Finally, residues that do not match the input sequence are pruned, producing *model_prune*, the default model used for assessment unless otherwise specified.

### Quality of the model building for 1-4 Å resolution maps

To benchmark CryFold, we first tested it against 177 maps of resolution better than 4 Å from ModelAngelo. The specific metrics for performance evaluation are introduced in Supplementary Information S1. For objective comparison, the ModelAngelo models for these maps were downloaded from the website given in its paper (2). For new maps, atomic models were generated locally using the version v0.2.3, which has higher accuracy to other versions in our tests.

While both CryFold and ModelAngelo perform well when the map resolution is better than 3 Å (Fig. 2A, indicated by points in the upper right corner), CryFold’s advantages become evident as the map resolution decreases below 3 Å. Notably, with a higher completeness rate, CryFold also generates more accurate structures, as indicated by Cα RMSD (0.310 Å vs 0.334 Å) and backbone RMSD (0.338 Å vs 0.375 Å) (Fig. 2E).

**Fig. 2.**
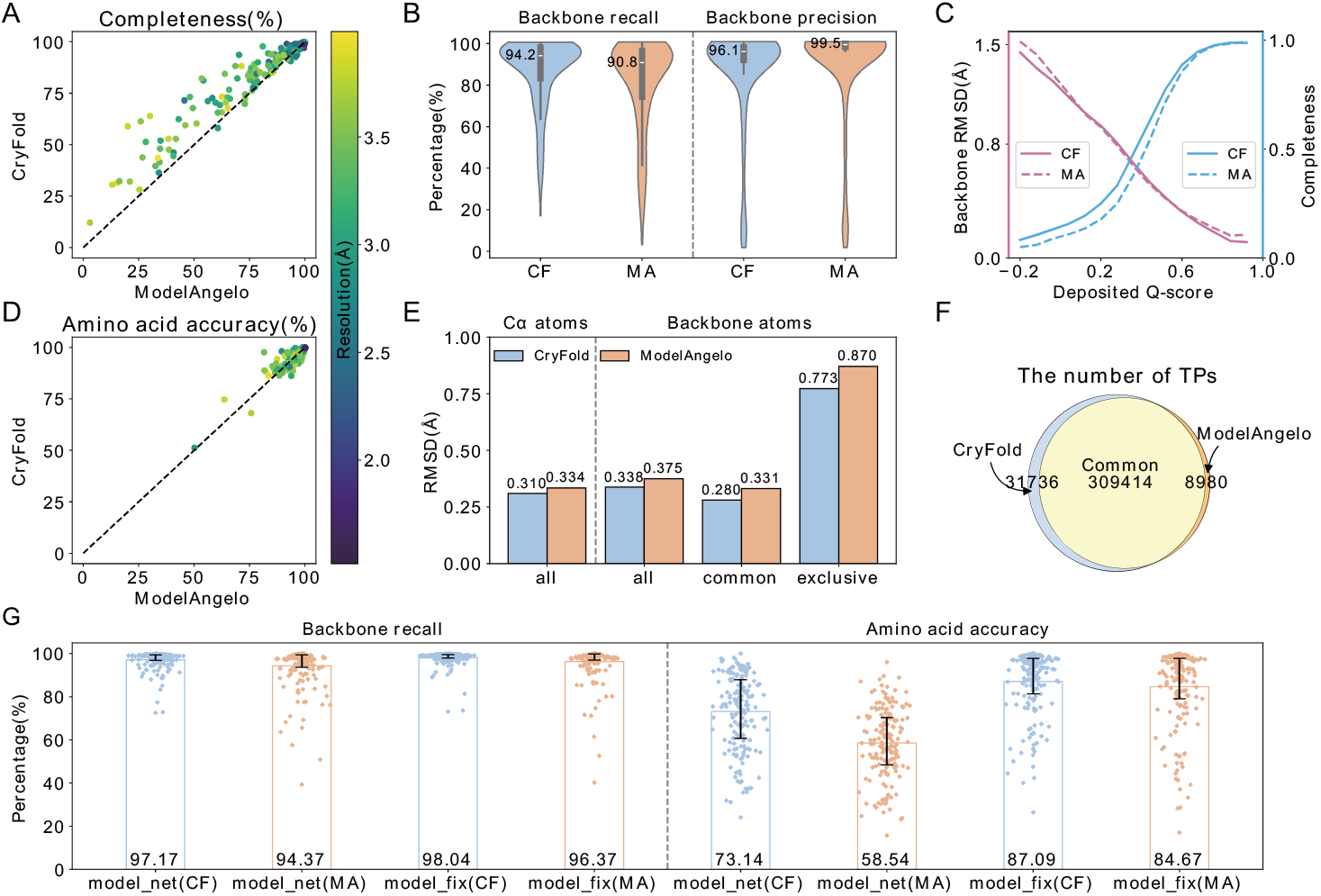
Comparison between CryFold and ModelAngelo on the test set of 177 high-resolution maps. (A) Head-to-head comparison based on the completeness. (B) Violin plots of the backbone recall and the backbone precision. (C) Backbone RMSD and completeness plotted as a function of the Q-scores of the deposited structures. (D) Head-to-head comparison of the amino acid accuracy. (E) Bar plots comparing the average RMSDs of Cα atoms and backbone atoms. The sizes of the sets of true positive residues (*all, common, exclusive*) are shown in (F). (F) Venn diagram showing the numbers of true positive residues predicted by both methods. (G) Backbone recall and amino acid accuracy for the intermediate models.

Completeness is closely linked to backbone recall and amino acid accuracy. A decline in either of these metrics results in reduced completeness. To further investigate this relationship, we calculated backbone recall and amino acid accuracy for the intermediate models (model_net and model_fix). CryFold achieves a backbone recall in model_net of 97.17% vs. 94.37% of ModelAngelo (Fig. 2G). If segments less than 4 residues are retained, this parameter increases by 0.87% and 2.0% for CryFold and ModelAngelo, respectively. The smaller increase for CryFold might reflect that the structures generated by our network are less fragmented.

Additionally, CryFold exhibits a higher amino acid accuracy before pruning (in both model_net and model_fix, Fig. 2G), and the two methods show comparable accuracy after pruning (in model_prune, Fig. 2D). The amino acid accuracy for the model generated by Cry-Net (model_net) is 73.14%, compared to 58.54% for ModelAngelo. This indicates that our network possesses a stronger ability to assign amino acid types, likely due to the enhanced transformer architecture. The HMM-based correction from ModelAngelo performs well, improving amino acid accuracy in model_fix to 87.09% for CryFold and 84.67% for ModelAngelo. In summary, CryFold shows high backbone recall and amino acid accuracy, contributing to the overall completeness.

We also analyzed the accuracy of predicted structures based on the Q-scores (38) of residues in the deposited structures. The Q-score reflects the consistency between the map density and the atomic structure, with higher values typically indicating better local resolution. Fig. 2C plots the backbone RMSD and the completeness of models built by CryFold and ModelAngelo against the deposited Q-scores. When the Q-score is less than 0 or greater than 0.6, CryFold models exhibit lower backbone RMSD. For other Q-score ranges, the RMSDs of both methods are comparable. In terms of completeness, CryFold outperforms ModelAngelo when the Q-score is below 0.6.

In addition to Q-scores, we also calculated the MolProbity scores (39) and EMRinger (40) metrics. The former reflects the geometric validity of the protein models, while the latter indicates the degree of agreement between the protein side chains and the density map. CryFold achieves better model quality than ModelAngelo while having a higher level of completeness (MolProbity scores 3.56 vs. 3.69, EMRinger 3.02 vs. 2.90). Detailed results can be found in Table S1.

Next, we compared the number of true positives (TPs) in the structures generated by both methods. A residue is classified as a TP if a corresponding deposited residue exists within 3 Å. Both methods have 309,401 common TPs, with backbone RMSD values of 0.28 Å and 0.33 Å for CryFold and ModelAngelo, respectively (Fig. 2F). Additionally, CryFold identifies 31,736 TPs that ModelAngelo misses, contributing to its higher completeness. These residues generally correspond to lower-resolution areas, as indicated by their lower Q-scores (0.47) and higher backbone RMSD (0.77 Å, Fig. 2E). In contrast, ModelAngelo identifies only 8,980 TPs (with a backbone RMSD of 0.87 Å) that are missed by CryFold. This data reinforces CryFold’s advantage in constructing atomic structures for lower-resolution maps/regions.

### CryFold extends the map resolution limit for structure construction

Our tests indicate that CryFold exhibits slightly lower backbone precision compared to ModelAngelo (Fig. 2B), which may stem from CryFold’s ability to construct structures in lower-resolution regions (see below), which are often absent in deposited structures and classified as false positives during evaluation. For instance, in the cryo-EM structure of the *type III-E CRISPR Craspase gRAMP-crRNA complex* (PDB ID: 7Y82, EMD-33678; green cartoon and gray surface in Fig. 3A), residues 1041-1380 are largely absent in both the deposited structure (green cartoon in Fig. 3A) and the ModelAngelo model (magenta cartoon in Fig. 3A). The structures for these regions are modeled by CryFold (blue cartoon in Fig. 3A, B) but classified as false positives, resulting in a lower precision than ModelAngelo (88.8% vs 99.8%). However, these residues are modeled with high confidence (>70, see the next section). We also submitted the domain sequence to AlphaFold3 (41) (AF3) for structure prediction, yielding a high confidence score (AF3 confidence score > 0.8). Structure superimposition indicates that the AF3 model closely resembles the CryFold model (TM-score 0.86, RMSD 2.07 Å; Fig. 3C). This cross-validation supports the correctness of the CryFold model for the missing residues.

**Fig. 3.**
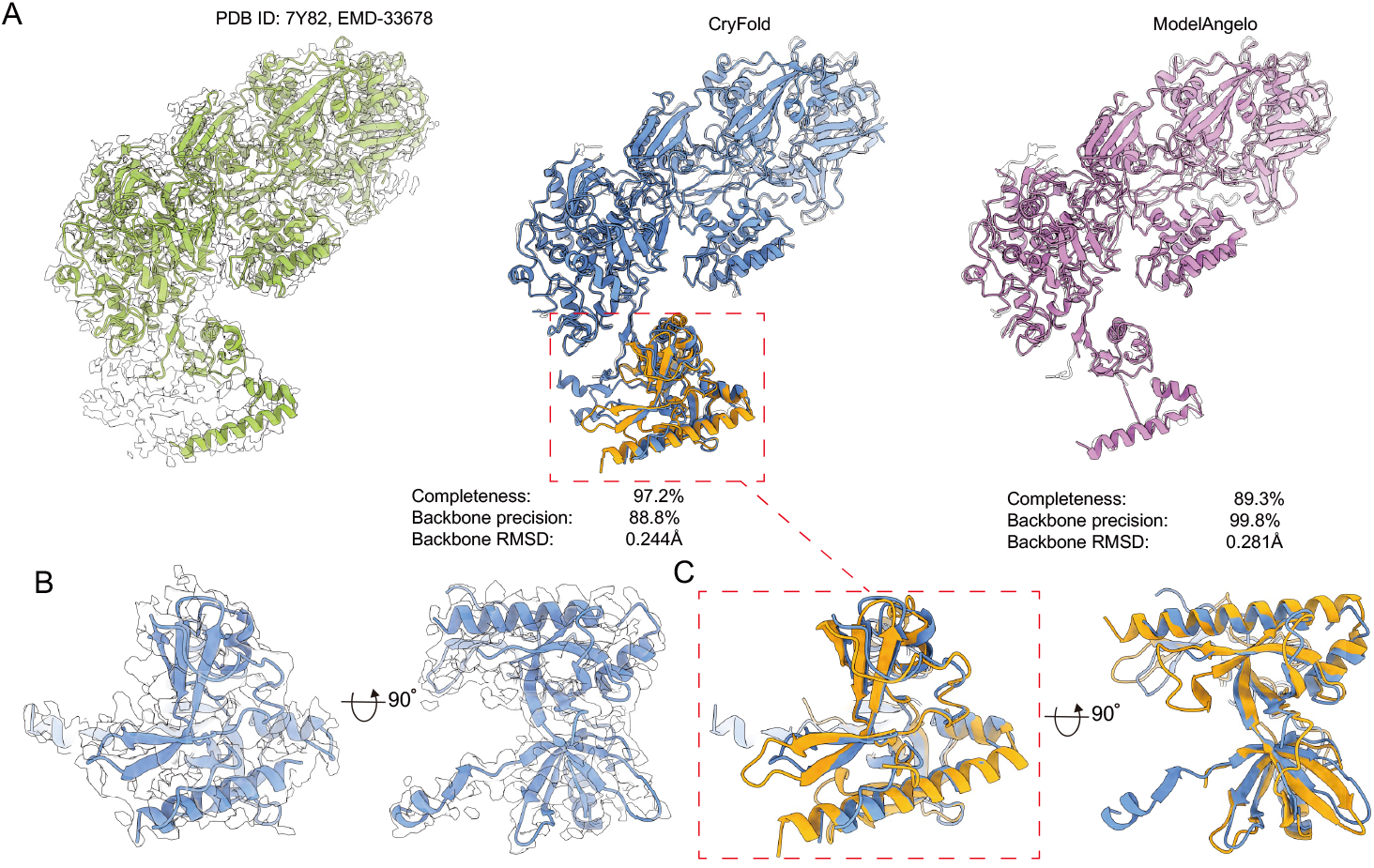
An example of how CryFold has a more complete model compared to the deposited structure. (A) The gray surface is the density map EMD-33678 (reported resolution 2.83Å). The green/gray cartoon is the deposited structure (PDB ID: 7Y82); the blue cartoon is the CryFold model, the magenta cartoon is the ModelAngelo model, and the orange cartoon is the AF3 model for the unmodeled regions in the deposited structure. (B) The blue cartoon is the modeling of the absent residues 1041-1380 in the deposited structure by CryFold and the gray surface is the corresponding density map. (C) An enlarged view of the red box in (A).

The above phenomenon is not isolated, and we have observed it in numerous cases, potentially impacting overall backbone precision. On average, the CryFold models for the 177 maps include approximately 75,000 new residues predicted with high confidence (> 60, see Fig. S1) absent in deposited structures. All structures generated by CryFold are available for download at https://yanglab.qd.sdu.edu.cn/CryFold.

### Evaluation of residue confidence scores

CryFold provides a reliable estimation of model accuracy through a per-residue confidence score, similar to the confidence scores available in protein structure prediction. This score is derived from the predicted FAPE loss (see Methods). The confidence score ranges from 0 to 100; a higher score indicating lower FAPE loss and a more confident prediction. This score is stored in the B-factor field of the mmCIF file. We analyzed the correlation between the confidence scores of all residues in 177 predicted structures generated by CryFold and their backbone RMSDs (Fig. 4A). Generally, a higher residue confidence score correlates with a lower backbone RMSD. In Fig. 4B and 4C, we illustrate two examples colored by their respective confidence scores. Notably, CryFold assigns lower confidence scores to loop regions, which tend to be more flexible and exhibit lower resolution in density maps.

**Fig. 4.**
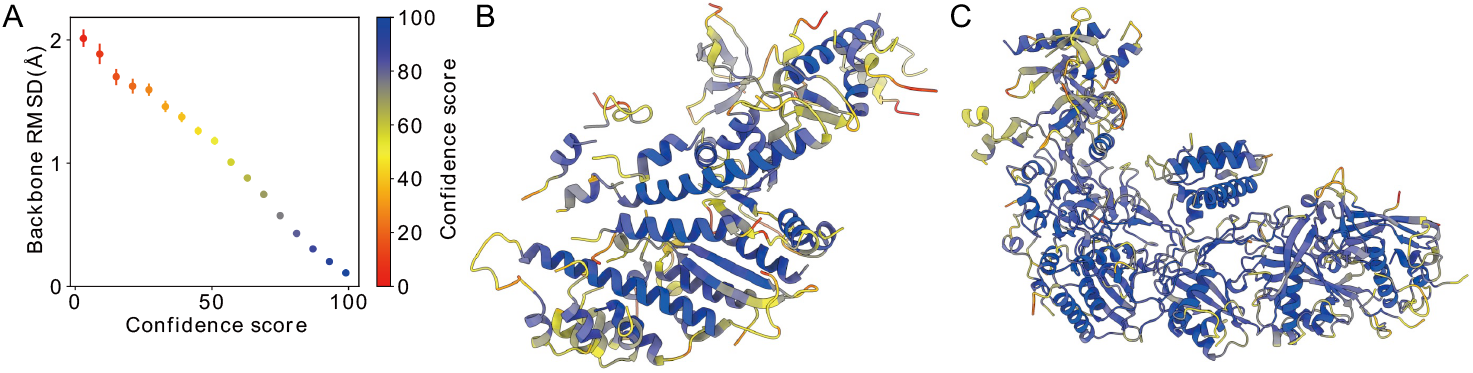
Correlation analysis between CryFold confidence score and backbone RMSD. (A) Confidence score bin size is 6, and the error bars represent the 99% confidence interval of the mean on a per-residue basis. (B-C) CryFold models colored by confidence score for the maps EMD-28080 (b, PDB ID: 8EFD, reported resolution 3. 8Å) and EMD-33678 (c, PDB ID: 7Y82, reported resolution 2.83 Å).

### Map masking improves backbone precision

Another factor contributing to CryFold’s lower precision is the absence of automatic map masking. Map masking is typically used to exclude regions of low resolution or those unrelated to the structures of interest (e.g., membrane or solvent). We evaluated the impact of map masking on a set of 70 maps from the 177 high-resolution maps, for which both the original and masked maps are available from EMDB (42). The average backbone precision for structures built by CryFold increased from 86.0% (using the original map) to 90.7% (using the masked map). While most cases show no significant difference between the original and the masked maps (Fig. S2), some examples demonstrate improved backbone precision with masked maps. For instance, for the map EMD-33306, the backbone precision improved from 25.2% to 97.1% after masking (Fig. S2E). However, map masking can sometimes result in incomplete atomic structures (Fig. S2F). Nonetheless, our package provides an optional argument to accept masked maps for expert users.

### Benchmarking performance at 4-7 Å resolution maps

Next, we assessed the performance of CryFold at lower resolution range. We tested 104 maps with resolutions of 4-7 Å (Fig. 5A-C). As expected, the accuracy is lower compared to high-resolution maps. Nevertheless, CryFold builds relatively complete models (>50% completeness) for approximately 40% of the tested maps. CryFold generally constructs more complete models than ModelAngelo (backbone recall 43.0% vs. 25.8%; completeness 36.8% vs 23.6%) while maintaining similar accuracy (amino acid accuracy 86.6% vs. 85.3%; backbone RMSD 0.777 Å vs 0.905 Å). An example is given in Fig. 6A to illustrate the performance on low-resolution maps.

**Fig. 5.**
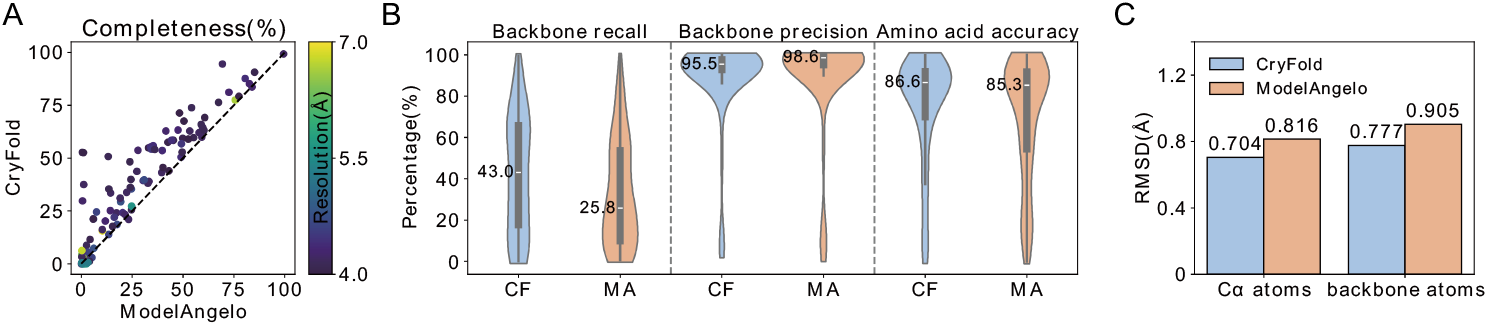
Comparison of CryFold and ModelAngelo on 104 test maps with resolutions between 4-7 Å. (A) Head-to-head comparison of the completeness. (B) Violin plots of backbone recall, backbone precision, and amino acid accuracy. (C) Bar plot of the main-chain atom RMSD.

**Fig. 6.**
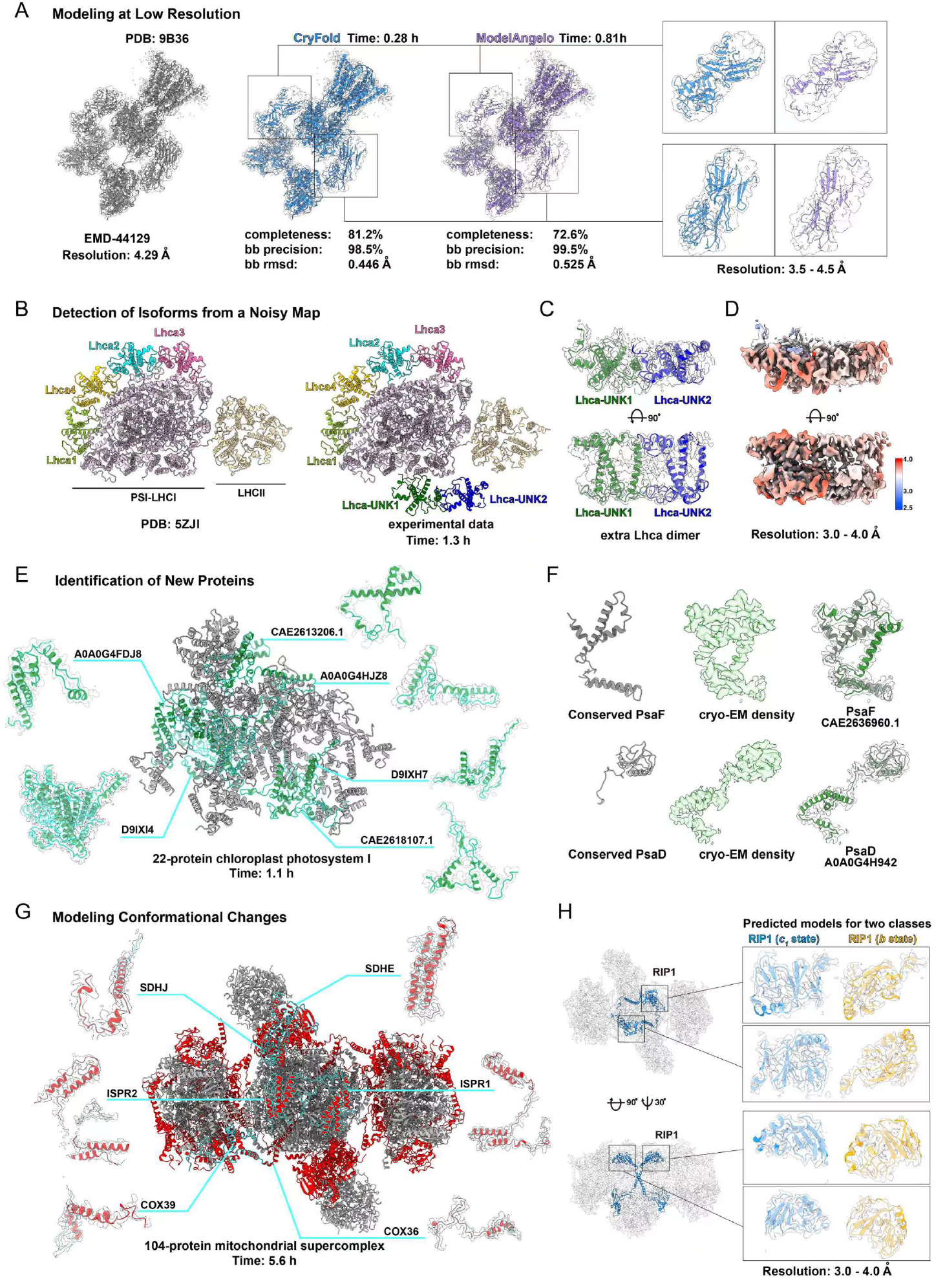
Capabilities of CryFold with four complementary examples. (A) Modelling at low-resolution regions showing comparison between the deposited model PDB ID: 9B36 (grey cartoon, left), the CryFold model (blue cartoon, middle), and the ModelAngelo model (purple cartoon, right) with corresponding closeup views. The density map EMD-44129 (reported resolution 4.29 Å) is shown in transparent. (B) Detection of isoforms from a noisy map is shown with a plant Photosystem I, where CryFold modelled two additional proteins (right) compared to the maize complex (left, PDB: 5ZJI). (C) CryFold identified two specific isoforms in the noisy map region with 3.0-4.0 Å resolution (transparent surface) from ten candidates with over 75% sequence identity. (D) The same map region colored by local resolution, showing the ability of CryFold to segment the density, as a major fraction is contributed by various cofactors. (E) Identification of six new proteins (green) in a 22-protein photosystem complex, shown with their corresponding map densities. (F) Two substantially extended proteins identified by CryFold in the same complex (right) compared to the canonical models (left). (G) The CryFold model of the mitochondrial respirasome II2-III_2_ -IV2 (16) (grey) with 52 out of 104 proteins that were not reported in other complexes (red). Six of these proteins are shown with their densities and compared with the ModelAngelo in Fig. S10. (H) Two distinct conformations (blue and yellow in close-up views) of a single protein (RIP1) in the 104-protein supercomplex modelled by CryFold in two classes.

### Performance on a non-redundant set of maps

The above test maps were collected based on the release date (i.e., after the training maps), without control of sequence redundancy to the training maps. To investigate this issue, the 281 test maps (from both the high-resolution and the low-resolution test sets) are compared with the training maps at a 40% sequence identity threshold, resulting a non-redundant set of 101 maps (75 and 26 from the high-resolution and the low-resolution test sets, respectively).

The performance of CryFold is listed in Table S4. Surprisingly, the CryFold models for the non-redundant maps are more complete than the redundant maps (completeness 74% vs. 66%). By dividing these maps into two subsets according to their resolutions, we can see that sequence redundancy does not affect the high-resolution maps (completeness 83% vs. 84%). However, the model completeness for the non-redundant low-resolution maps is higher than the full test set (50% vs. 37%). This is because, before removing redundancy, there were 32 poor models in the low-resolution dataset, where the completeness of the CryFold models were below 20%. Only three such models were retained in the non-redundant set, resulting in higher completeness. In summary, these results indicate that the key factor impacting CryFold’s performance is the map resolution rather than the sequence similarity to the training data.

### Model accuracy in Stage 1 and Stage 2

CryFold works in two stages (see Fig. 1), predicting Cα atoms in Stage 1 and all atoms in Stage 2. We compare the results of CryFold and ModelAngelo in the Stage 1 predictions (see Fig. S3B-D Stage 1). CryFold shows higher recall than ModelAngelo at the expense of lower precision and higher RMSD (backbone recall 99.0% vs. 98.4%; backbone precision 75.5% vs. 90.4%; Cα RMSD 0.729 Å vs. 0.640 Å). This is because CryFold returns much more number of candidate Cα atoms than ModelAngelo (on average 8770 vs. 4677 per map), resulting lower precision and higher RMSD at this stage. However, most false Cα atoms are successfully filtered out by the network Cry-Net at Stage 2, which can be seen from increased precision from 75.5% to 95.8%.

As shown in Fig. S3E-G, we compare the MolProbity scores (39) and EMRinger (40) of atomic models constructed by CryFold and ModelAngelo. CryFold achieves slightly better model quality than ModelAngelo (MolProbity scores 3.66 vs. 3.76, EMRinger 2.45 vs. 2.40). The MolProbity scores for both methods are not ideal, which may be related to the models constructed by deep learning methods not undergoing a relaxation step. Here, structures with MolProbity scores >4 (about 21 models) were further relaxed using the refine module in trRosetta (35), resulting in new models with improved MolProbity scores to 2.1 (see Fig. S3F).

Finally, we compare CryFold and ModelAngelo on the intermediate models (model_net) without any post-processing. The results (see Fig. S3H) show that CryFold has a higher backbone recall (95.5% vs. 89.7%) and higher amino acid accuracy (65.9% vs. 51.3%). This demonstrates the superiority of the Cry-Net in constructing atom positions and recognizing amino acid types.

### Comparison with other methods that use additional constraints

We further compare CryFold with two recent methods that incorporate additional constraints, including CryoFold (43) and DeepMainmast (28). CryoFold is a method for predicting protein complex structures, utilizing density maps, multiple sequence alignments (MSAs), and homologous structure templates. DeepMainmast constructs complex structures from density maps by deep learning and predicted structures from AF2.

At the time of this study, CryoFold was not open-source; however, a web server was available at https://cryonet.ai/cryofold. We manually submitted density maps and protein sequences to the CryoFold web server for modeling, which took an average of approximately 2 hours per target in our tests. To streamline testing, we randomly selected 25 density maps from the above non-redundant dataset, spanning resolutions of 2.2-4.2 Å, for comparison with CryoFold.

CryFold demonstrates competitive performance relative to CryoFold (average completeness: 80.8% vs. 81.5%; amino acid accuracy: 93.1% vs. 91.7%; backbone RMSD: 0.406 Å vs. 0.436 Å). Owing to additional constraints from MSAs and templates (it remains unknown if homologous templates were excluded or not), CryoFold slightly outperforms in completeness for most samples (Fig. S4A). However, CryFold exhibits greater stability in completeness, with a lower standard deviation (22.3% vs. 25.0%).

The DeepMainmast protocol comprises two components. The first, DeepMainmast(base), constructs models using *de novo* main-chain tracing guided by deep learning. The second integrates the DeepMainmast(base) model with structure predictions from AF2. Modeling a target with DeepMainmast typically requires several hours. For convenience, we referenced the data from the original DeepMainmast paper (28), which includes results for 20 protein complexes. To eliminate bias from similar targets in our training dataset, we excluded complexes sharing over 40% sequence identity with the training set, yielding a final set of 17 protein complexes. The results of the other three protein complexes are highlighted in blue in Table S8. Additionally, we found that using the original density map (EMD-5495) directly as input for CryFold yields very poor results. This may be related to the fact that EMD-5495 was released early (in 2012) with high background noises. So we preprocessed EMD-5495 in the same way as ModelAngelo and then used it as input for CryFold.

We first compare CryFold with DeepMainmast(base), which a fair comparison given their identical input information. Fig. S5 indicates that CryFold outperforms DeepMainmast(base) for almost all complexes (completeness: 83% vs. 64%; amino acid accuracy: 93% vs. 83%; TM-score: 0.77 vs. 0.63). When compared to the full DeepMainmast protocol, CryFold remains competitive (completeness: 83% vs. 87%; amino acid accuracy: 93% vs. 96%; TM-score: 0.77 vs. 0.92).

Note that TM-score may not be an appropriate metric for evaluating CryFold, as it relies on density maps and protein sequences for structure construction, which can lead to incomplete predictions for map regions with poor resolution. In contrast, methods like AF2 generate full-length models for all residues in a complex, regardless of map resolution.

To summarize, the comparisons above demonstrate that CryFold remains competitive with CryoFold and DeepMainmast, despite the additional constraints utilized by these methods. We intentionally avoided incorporating additional constraints into CryFold to prioritize ease of use and efficiency. As a result, CryFold is expected to be several times faster than both methods. Nevertheless, enhancing model building for density maps with poor resolution, potentially by integrating predicted structure models, could be beneficial. We defer such improvements to future developments of CryFold.

### Sensitivity analysis of CryFold to hyperparameters

CryFold has many hyperparameters during the inference phase. Here, we analyze two key hyperparameters: the prediction threshold for the classification network in Stage 1 (denoted as *t*) and the number of recycling rounds in Stage 2 (denoted as *n*). In the default settings of CryFold, *t* is set to 0.6 and *n* to 3. We performed tests to evaluate the effects of varying these parameters.

As illustrated in Fig. S6A, reducing *t* and *n* affects the completeness, with *n* having a stronger impact on the completeness. This suggests that recycling effectively enhances the model completeness. Additionally, when lowering the Stage 1 threshold to *t* = 0.4, the backbone precision remains largely unaffected (see Fig. S6B), indicating that CryFold effectively filters out false positives generated in Stage 1.

We also tested the impact of randomly rotating and translating density maps. We fed the transformed density map into CryFold. Then, the models output by CryFold are aligned with the native models using US-align (44) for comparison. The results were interesting: appropriate rotation and translation can improve model quality (see Fig. S6), although models built from the original and the transformed maps remain broadly similar. These data demonstrate the robustness of CryFold.

### Ablating CryFold components to assess drivers of performance

First, we checked how much improvement is introduced by the *L*_2_ regularization with simulated density maps (i.e., in Eq. 1). For the convenience of the experiment, we arranged three configurations of U-Net for study: the default version (CryFold Stage 1), CF with *L*_2_ loss (CryFold Stage 1 without a pre-training process, see Supplementary Information S2.2), and CF w/o *L*_2_ loss (neither a pre-training process nor *L*_2_ loss constraints). Fig. 7A, and 7B show that the models at Stage 1 become worse after removing the pre-training process (backbone recall 99.0% vs. 97.0%; backbone precision 75.5% vs. 70.5%; Cα RMSD 0.729 Å vs. 0.852 Å). Removing the *L*_2_ regularization has a slight impact on performance (backbone recall 97.0% vs. 96.3%; backbone precision 70.5% vs. 71.2%; Cα RMSD 0.852 Å vs. 0.854 Å).

**Fig. 7.**
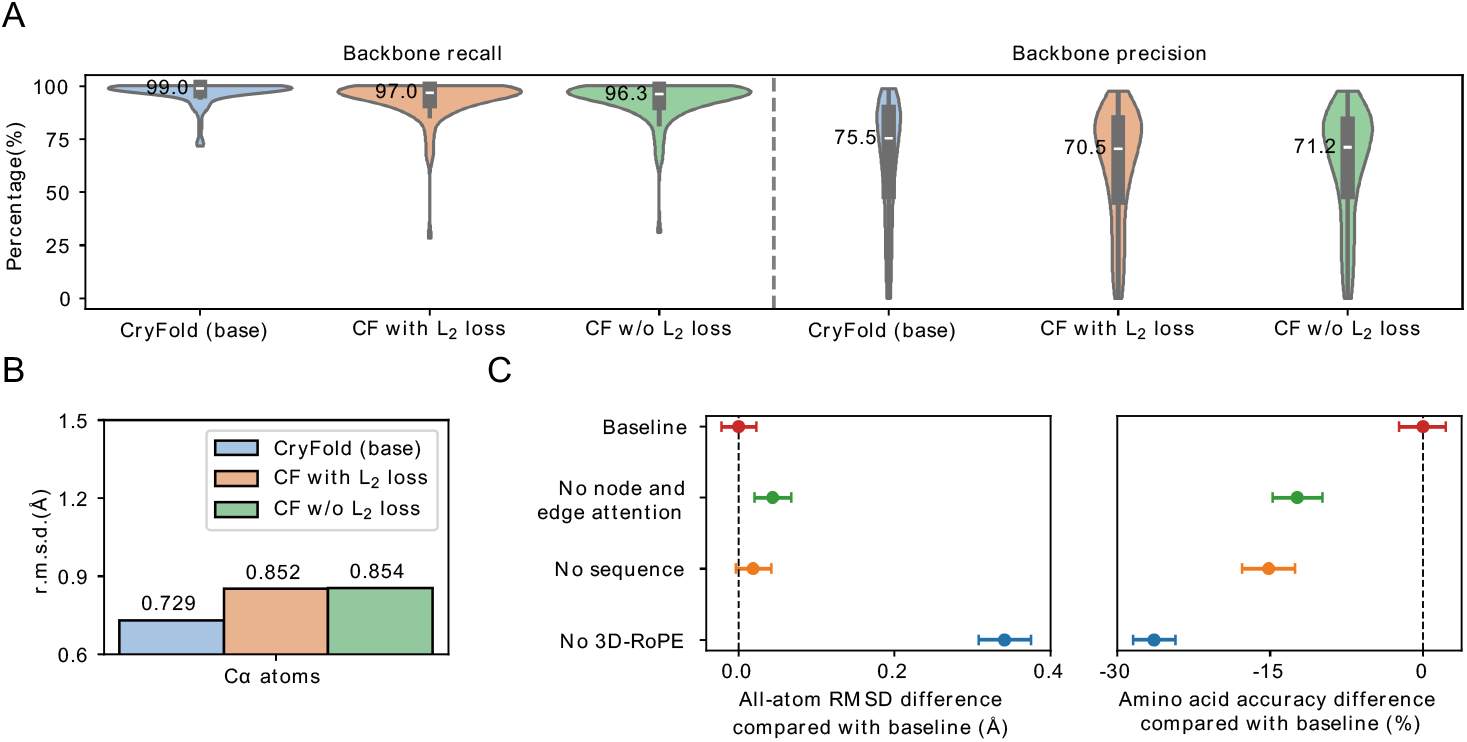
Results of ablation study. The experiments for (A-B) are conducted on the non-redundant test set. We mainly studied CryFold Stage 1 under three configurations. (A) is a violin plot regarding backbone recall/precision. (B) is a bar chart for the Cα RMSD. (C) The experiments on 177 high-resolution maps. Left is the all-atom RMSD. Right is the amino acid accuracy. The mean is indicated by the center point. The error bar is calculated based on the 95% percentile interval.

To investigate the contributions of different components to CryFold, we conducted ablation experiments on the 177 high-resolution maps. Four configurations of its network, Cry-Net, were assessed: the default version (18-layer Cryformer), a version without node and edge attention, a version without sequence input, and a version without 3D rotary position embedding (3D-RoPE). These experiments evaluate the impact of network depth, sequence information, and 3D-RoPE on performance. Metrics included amino acid accuracy and all-atom RMSD, calculated against the deposited structures (Fig. 7C). The default model achieved 0.506 Å all-atom RMSD and 73.1% amino acid accuracy. The RMSD increased slightly for shallower networks and the version without sequence input, while amino acid accuracy dropped significantly, especially for the version lacking sequence input. The configuration without 3D-RoPE exhibited the most substantial differences, with an increased all-atom RMSD of 0.341 Å and a 26.4% decrease in amino acid accuracy. Notably, the loss curve for the ablation experiment without 3D-RoPE displayed oscillations during training (Fig. S7), suggesting that this encoding is crucial for the effective training of Cry-Net.

### Running time and GPU memory usage

We evaluated CryFold’s running time as a function of protein length on the 177 high-resolution maps using a single GPU (A100, ∼15G memory) (Fig. S8A). The running time increases linearly with protein length. CryFold can build a structure of approximately 40,000 residues in about 3.5 hours. CryFold allows users to balance GPU memory and running time. By default, CryFold processes 300 residues at a time, using ∼13 GB of GPU memory. An additional ∼1 GB of memory is required for every 100 extra residues. Fig. S8B illustrates the running time for the PDB entry 8FNV (>9,000 residues) under three configurations. Increasing GPU memory by 1.6 times (from 15GB to 24GB) results in acceleration of the speed by 1.5 times (from 44 minutes to 30 minutes). This linear relationship can be generalized to proteins of any length. In additional tests, the accuracy of predicted models remains consistent with increased crop size.

### Case studies to assess model completeness, protein identification and conformational changes

We showcase the advanced capabilities of CryFold with four complementary examples, each addressing a distinct and challenging aspect of modern macromolecular modeling (Fig. 6). They are introduced below in details.

#### 1) Handling low-resolution regions within maps

An illustrative low-resolution example is provided in Fig. 6A, comparing models generated by CryFold and ModelAngelo. The case involves an open state of *kainate receptor GluK2* (PDB ID: 9B36) with 4.3 Å resolution. CryFold successfully constructs structures for 81.2% of the residues, achieving 0.45 Å backbone RMSD. In contrast, ModelAngelo constructs structures for 72.6% residues, achieving 0.53 Å backbone RMSD. Due to the lower resolution in the peripheral regions of the density map, ModelAngelo encountered difficulties during modeling. However, CryFold constructed relatively complete atomic models for these regions (Fig. 6A). In addition, CryFold takes less time than ModelAngelo for the modeling (0.28h vs 0.81h). This example clearly demonstrates CryFold’s superior modeling capability and efficiency for handling low-resolution regions within maps.

#### 2) Resolving compositional heterogeneity by identifying closely related isoforms within a noisy map

To illustrate CryFold’s precision in distinguishing closely related sequences, we selected a challenging case from photosynthesis involving light harvesting. This process is mediated by pigment-protein light-harvesting complexes (Lhc) which function as antennas around catalytic reaction centers, converting photons into chemical energy (45). These Lhc proteins have been conserved across eukaryotic photosynthetic organisms for billions of years (46). In the structure of a plant photosystem, the antenna consists of four Lhca proteins (47, 48). However, wild-type plants have been reported to contain additional isoforms with higher energy chlorophylls (49), and studies suggest extra Lhca dimers (50), supported by *in vitro* reconstitution (51, 52). We purified photosystem I from a marine angiosperm in the Mediterranean and reconstructed its 3D structure. No reference protein database was used for this modeling. Compared to canonical models from pea and maize (53, 54), we observed an additional Lhca regulatory dimer with unknown sequences (Fig. 6B).

These unidentified chains are located in a region with 3.0-4.0 Å resolution (Fig. 6C). Despite the short length (∼200 residues) and high similarity (>75% identity) with other isoforms, CryFold generated an atomic model for these new chains without any reference protein database. This enabled us to match the sequences within genomic data and identify them among a family of over ten highly similar proteins. Specifically, these chains were identified as Lhca1 and Lhca4 isoforms. Furthermore, this performance also underscores CryFold’s ability to segment cryo-EM maps, focusing only on protein regions even with multiple cofactors present, as well as noisy density (Fig. 6C,D). Specifically, the map region for the two newly identified proteins includes more than 40 non-protein cofactors such as chlorophyll a, chlorophyll b, lutein, violaxanthin, and β-carotene, as well as various lipids distributed on different sides of the heterodimer that dominate the cryo-EM density (Fig. 6D). The calculation took 1.3 hours on an A800 GPU.

To assess the quality of the assignments, we used the sequences of the newly identified Lhca-UNK1 and Lhca-UNK2 proteins, along with their interacting chains, as input to AF3 (41). The AF3 model is relatively accurate (ranking score: 0.71) and has an overall fold consistent with CryFold’s model (Fig. S9). However, the positions of the newly identified proteins are shifted, probably because the predicted interfaces with other interacting chains in the AF3 model have low accuracy (pTM: 0.37 and 0.3). This experiment illustrates the superior capability of CryFold over pure structure modeling.

#### 3) Detecting new proteins in multi-component complexes

Another challenge in protein modeling for complex organisms is determining the identity of previously uncharacterized proteins directly from the cryo-EM map. CryFold is able to address this challenge iteratively without extra model training. Specifically, an initial set of sequences, which can be completely random or part of the sequences related to the density map, can be fed into CryFold. The HMM profiles generated by CryFold are then compared with the sequences in the sequence database related to the density map (e.g., can be easily download from the UniProt database by organism information, or just use the whole UniProt database if nothing is known). Successfully matched sequences are added into the initial sequence set. CryFold is run again with the updated set of sequences. This iterative process continues until no new sequences are identified. The final set of sequences represents all sequences related to the density map. Once the sequence database is prepared, this process becomes fully automated and requires no manual intervention. This enables for the automatic search of sequences within the density map from a vast sequence database, thereby constructing atomic models. The prerequisite for this process to work is that the provided sequence database must encompass all protein sequences in the density map. Given the rapid development of the genome sequencing projects, this assumption is not difficult to meet. We can use this function to identify unknown or missing subunits that human experts failed to build.

We tested CryFold’s ability to tackle this using an unpublished reconstruction of another photosynthetic complex with multiple chains from a brown alga. In this example, CryFold identified and modeled 22 proteins, six of which had not been previously characterized (Fig. 6E). Additionally, two universally conserved chains exhibited fully resolved extensions of 71 and 99 residues, effectively doubling the original size of each protein in the core complex; indicating potential functional relevance (Fig. 6F). None of the newly modeled sections had not been identified before, nor had the new chains been predicted to be part of the complex. CryFold’s ability to detect new proteins and structural extensions within intricate cryo-EM data demonstrates its efficiency, with calculations taking only 1.1 hours on an A800 GPU.

#### 4) Capturing minor conformational changes in a mega-Dalton complex

To explore CryFold’s sensitivity to subtle conformational changes, we used a mitochondrial supercomplex functioning as a respirasome, with a mass of 1.8 MDa, in which only one protein displayed a slight rotation (16). First, CryFold generated an atomic model from the map of the native source, constructing the 104-protein complex in 5.6 hours, with half of the chains being species-specific (Fig. 6G). Compared to ModelAngelo, CryFold provided higher completeness in lower resolution regions of the map (Fig. S10). To further assess CryFold’s precision, we classified two maps and found a subtle 15 Å movement in a single peripheral domain of a protein. This minor motion was resolved in two conformations within the 261 × 296 Å supercomplex. The resolution in the area of this conformational shift is 3.0-4.0 Å. CryFold’s ability to capture such minor structural variations demonstrates its effectiveness for investigating dynamic biological particles and targeting structural regulation.

#### 5) Performance on a large protein-nucleic acid complex

Finally, we tested if CryFold works for large-scale biological macromolecules. As a representative example, we used the human mitoribosome, which has over 80 protein chains and a large overall dimensions of 307 × 287 × 286 Å ^51^. Although CryFold does not currently model nucleic acids, it successfully segmented the heterogeneous density, where one-third corresponds to nucleic acids ^52^. Notably, 78 out of 87 protein chains were modeled with >80% completeness (Fig. S11). This example demonstrates that CryFold effectively handles large-scale structures containing significant amounts of nucleic acids.

## Discussion

In this study, we demonstrate the advanced capabilities of CryFold in addressing complex challenges in macromolecular modeling through a novel local-attention network and 3D rotary position embedding. CryFold increases the accuracy and completeness of automated built protein models while reducing the execution time and requirement for map resolution. Tests on large complexes show identification of previously unknown proteins that could not be revealed by other automated tools or predicted. For most tested maps, it constructs models with over 80% completeness, including for protein complexes with nucleic acids, and previously missing residues could be modeled. Through specific examples, we illustrate CryFold’s applications in addressing challenging tasks of modern macromolecular modeling. They include: successfully resolving low-resolution map regions, identifying closely related isoforms within noisy maps, detecting novel proteins in multi-component complexes even without a well-established protein database, and capturing subtle conformational changes within large protein assemblies. CryFold also goes beyond simple structural modeling by segmenting maps with multiple cofactors. Together, these results underscore CryFold’s ability to tackle a variety of complex challenges in macromolecular modeling in a more time-efficient manner. The versatility, cost-efficiency and precision, mark CryFold as a valuable tool for high-throughput macromolecular modeling and functional analysis of dynamic biological assemblies.

Although CryFold leverages deep learning to reduce resolution requirements, modeling at low resolution remains challenging. This difficulty stems from the inability to accurately identify protein side chains in low-resolution regions. The challenge arises from the heterogeneous nature of cryo-EM data: stable residues are well-resolved (reflected in high-resolution areas), whereas dynamic regions appear blurred (reflected in low-resolution areas). One potential solution for low-resolution regions is to represent them with an ensemble of conformations rather than a single fixed structure. CryoFold (55) serves as a good example of using molecular dynamics simulations to address this issue. Additionally, CryFold can incorporate MDFF (56) during the post-processing stage, which is a method for flexible fitting of density maps using molecular dynamics. These enhancements will be explored in future work.

## Methods

### Training and test data

#### Training set and validation set

In order to train our method and compare with ModelAngelo(2), we constructed the training dataset with the same setting used in ModelAngelo. Specifically, the training set consists of 6422 EMDB (42) cryo-EM density maps released before April 1, 2022 with resolution better than 4Å and paired structures in PDB (6). The full training set was used to train an initial model. The model was used to filter the training dataset based on the precision (>0.5) of the predicted Cα atoms. After filtering, there are 5731 remaining map-model pairs, of which 10% are randomly selected for validation and 90% for training.

#### High-resolution test set

The test dataset we used consists of 177 high-resolution (better than 4 Å) maps from ModelAngelo(2) (released between April 1, 2022 and February 9, 2023).

#### Low-resolution test set

To test the performance of CryFold on low-resolution maps, we collected another set of maps with lower resolution. We first obtained 135 maps with resolutions 4-7 Å, which were released between January 1, 2024 and June 1, 2024 (about two years after the training data). Among these maps, ModelAngelo failed to build atomic structures for 31 maps (the program did not respond for a long time or encountered errors during the process). Therefore, the final low-resolution test dataset consists of 104 maps with resolutions 4-7 Å. The number of residues in these maps ranges from 250 to 17520 (3461 residues on average).

#### Non-redundant test set

To investigate if there is any issue of data leakeage or not, we construct a non-redundant set here. Specifically, if the test density map and the training density map have at least one pair of protein chains with over 40% sequence identity, we will remove it from the test dataset. Additionally, the test dataset is clustered internally based on 40% sequence identity. After applying the above steps, out of 281 map-model pairs, 101 map-model pairs are retained.

#### CryFold algorithm

CryFold builds the atomic structure from the cryo-EM density map and amino acid sequence information through two major steps (see Fig. 1). The first step predicts the Cα atom positions using U-Net(36); The second step builds all-atom positions using an enhanced transformer network Cry-Net. They are described in details below.

#### Step 1: Predict the Cα atom positions using U-Net

We train a 3D convolutional neural network to predict the Cα atom positions of all residues in the density map, which are represented by the Cα atoms coordinates. The convolutional neural network adopts the U-Net architecture(36). A detailed architecture can be found in the supporting Fig. S12. The input to the network is the cropped density map, with each box size being 64×64×64. The shapes of the network outputs (i.e., *y*^*pred*^ and *ŷ*^*pred*^ in Fig. 1) are the same as the original map but the value in each voxel (i.e., each cube 1×1×1) is the probability of containing a Cα atom (obtained through a sigmoid layer). The difference between *y*^*pred*^ and *ŷ*^*pred*^ is that their corresponding labels are slightly different; the label for the former is a hard label, while the label for the latter is a soft label. Here, *y*^*pred*^ is used as the final output of the U-Net.

The loss function for U-Net(36) consists of the focal loss (57) (*FC*) and *L*_2_ distance loss (*L*_2_) as follows:

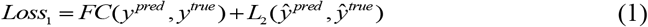

where *y*^*pred*^ and *ŷ*^*pred*^ are both predicted probability density maps for Cα atoms. *y*^*true*^ represents the map labeled by the positions of the real Cα atoms, in which a value of 1/0 indicates the presence/absence of a Cα atom; *ŷ*^*true*^ represents the simulated Cα probability density map, which is calculated the deposited Cα atom coordinates as follows(58):

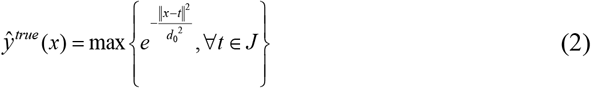

where *x* is a grid point, *t* represents the coordinates of the Cα atoms, and *J* represents the set of all Cα atoms in the deposited structure; *d*_*0*_ represents a normalization factor (3 Å here). If the distance from the grid point *x* to the nearest Cα atom is *d*_*0*_, the probability value for the grid point will be *1/e*. We combine both losses, aiming for the network to learn rich information from the density map.

The Cα atom coordinates predicted above are refined through the mean-shift algorithm(26, 59). The initial Cα atom coordinates with a probability value above 0.6 are selected first. Then the coordinates for each of the selected Cα atom are updated as the weighted linear sum of its neighboring Cα atoms within 1.7Å (including itself). After this update, the neighboring Cα atoms are removed because the Van der Waals radius of a carbon atom is 1.7Å. Such refinement is necessary because the real coordinates are continuous while the predicted ones from the density maps are discrete. After the refinement, we obtain a set of predicted Cα atoms (the total number of Cα atoms is denoted by *r*).

#### Step 2: Build all-atom positions using Cry-Net

Inspired by AlpahFold2, we designed a transformer network to build all-atom positions using a transformer network (named Cry-Net) from the input density map, the input sequences and the predicted Cα atoms. Cry-Net consists of two transformer-based modules. The first one is called Cryformer (see the lower panel of Fig. 1, and Fig. S13A) and the second one is called Structure Module (see the lower panel of Fig. 1, and Fig. S14). Cry-Net iteratively updates two key representations of the protein structure: *node representation* and *edge representation* (correspond to the single representation and the pair representation in AF2, respectively), which are introduced below.

#### Node representation

The node representation (*r*×*c, r* is the number of predicted Cα atoms, *c* is the number of channels) is constructed from the predicted Cα atoms and the density map as follows. A cubic box with size 17×17×17 Å^3^ centered at each of the *r* predicted Cα atoms is cropped from the density map first. Then to extract information from the density map and reduce the dimensionality, a convolutional neural network (CNN) is used to convert the density map in each cubic box into *c* channels. The architecture of this CNN is the same as the encoder module in U-Net(36) (*i*.*e*., Fig. S12A).

#### Edge representation

The edge representation (*k*×*r*×*c*) is constructed similarly, measuring the relationship between neighboring nodes (*k* is the number of nearest neighbors considered). For each edge, a rectangular box with size 3×3×12 Å^3^ is cropped from the density map. The Cα atom is located at the center of the bottom box (*i*.*e*., the 3×3 square) and the direction of the box is the same as the edge vector. Finally, another convolutional neural network (with kernel size 3×3×1) is used to convert the density map in each rectangular box into *c* channels.

Note that the above representations are different with the ones used in AF2. The representations here are constructed from the density map guided by the predicted Cα atoms (inspired by ModelAngelo); rather than from multiple sequence alignment (MSA) and homologous templates in AF2. In addition, the edge representation (called pair representation in AF2) is a square matrix (*r*×*r*) in AF2; while a non-square matrix (*k*×*r*) here. This is because the density map already provides rich structure information and we only need to consider the *k*-nearest neighbors rather than all nodes.

The input and output of Cry-Net in the *n*-th iteration are defined by the following equation:

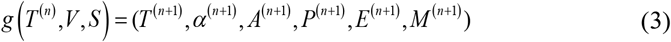

where *T* (^*n*^) ∈ ∼ ^*r*×3×4^ is the backbone frame (N, Cα, C) at the *n*-th step, *V* is the density map, *S* ∈ ∼ ^*r* ×1280^ is the sequence embedding from the protein language model ESM-2(60), *α* ∈ ∼ ^*r*×7×2^ is the sine and cosine encoding of the three backbone and up to four side-chain torsion angles, *A* ∈ ∼^*r×* 20^ is the probability vector for the 20 types of amino acids, *P ∈* ∼ ^*r*^is the confidence score of each node, *E* ∈ ∼ ^*r*×*k*^ is a binary vector indicating whether each node is sequentially adjacent to its *k* nearest neighbors or not, and *M* ∈ ∼ ^*r*^ is the probability of a node being a true residue.

#### Cryformer

The Cryformer module consists of a total of 18 blocks with non-shared weights, as shown in Fig. S13A. The key difference between Cryformer and the Evoformer in AF2 is the concentration on local attention rather than global attention. This is reflected by the number of neighbors considered (i.e., the parameter *k*). Evoformer will be equivalent to Cryformer when *k* equals to the total number of nodes (i.e., *k*=*r*). The key components of Cryformer are described below.

##### Sequence attention

This layer incorporates the protein sequence information into the node features with a gated cross-attention mechanism (see Supplementary Information S2.3, Algorithm S1). The node representations are queries, and the sequence embeddings from a pre-trained protein language model (*i*.*e*., ESM-2(60)) are keys and values.

##### Node attention

As shown in Fig. S13B, the node attention updates the representation of a given node based on self-attention on the node and its *k* nearest neighbors, with the information of edge representations serving as bias. As the depth of this network increases, it can collect information from more nodes, resulting in a larger receptive field. It is worth noting that this node attention will be equivalent to the row-wise gated self-attention in AF2 when *k* equals to *r*.

##### Transition and outer product

Inspired by AlphaFold3(41), The transition layer uses the SwiGLU(61) instead of ReLU(62) to update the node and edge features. It uses the non-linear activation function to enrich the representation of the two types of features. The outer product layer updates the edge features based on the node features. It incorporates the node features of two adjacent nodes into the edge features.

##### Edge attention

As shown in Fig. S13C, the edge attention updates the edge representation through self-attention mechanism. The information of edge *ij* is aggregated from the information of all edges connected to node *i* and all edges connected to node *j*. The edge attention can be seen as a special type of the node attention, according to the Whitney graph isomorphism theorem (Fig. S15). In addition, it is worth mentioning that when *k*=*r*, the edge attention is equivalent to applying AF2’s row and column attentions simultaneously.

##### 3D Rotary Position Embedding

(3D-RoPE). 3D-RoPE is not an independent component. Originally, RoPE(37) was used to process 1D sequences. We extend it to a node position encoding in 3D space. It is inserted into the node attention as a multiplicative positional encoding. It elegantly encodes the position information of each node into the attention calculations. The attention score will decay when the distance between two nodes increases. Please refer to the Supplementary Information S2.4 for more detailed description about 3D-RoPE.

##### Amino acid probability for each node

In order to generate all-atom positions, we try to assign each node to one of the 20 possible amino acids. This is done by using the information from the density map of each node (i.e., the original node representation), the information from neighboring nodes (i.e., the updated node representation) and the sequence information (obtained from the sequence attention in Cryformer). A separate MLP is trained to generate the amino acid probability vectors for all nodes (i.e., *A* in Eq. 3, Fig. S16). Ablation experiments indicate that removing the information of neighboring nodes (i.e., no node attention and edge attention) or removing the sequence information will affect the subsequent determination of amino acid identity.

#### Structure Module

The Structure Module here (see Fig. S14) is largely similar to the one in AF2 but with a few key differences (detailed below). It predicts the backbone frame (i.e., *T* in Eq. 3) as well as the backbone and side-chain torsion angles (i.e., *α* in Eq. 3), using the updated node and edge representations from Cryformer. The Structure Module consists of 7 blocks with shared weights. Two types of attention mechanisms are used in this module (see Supplementary Information S2.3 and Algorithm S2).

The first one is a self-attention mechanism applied to the node features (similar to the node attention in Cryformer). Given the constraints from the density map, the attention is restricted to each node and its *k* nearest neighbors (rather than all other nodes in AF2).

In addition, the position information between nodes is introduced into the attention mechanism using 3D-RoPE, ensuring that the attention score incorporates position information.

The second type of attention is the Invariant Point Attention (IPA) from AF2. It is conducted on the backbone frames and is invariant under global Euclidean transformations. Like the first type of attention, the attention is restricted to a node and its *k* neighboring nodes (rather than all others in AF2).

Cry-Net will be run iteratively to improve its output. During training, 0-2 rounds of iterations are performed. Only the backbone frame is iterated. During inference, two rounds of iterations are performed. The reason for choosing *n*=2 here is to keep a balance between computational cost and performance. No significant improvement in the network output was observed when more rounds of iterations are performed.

Finally, all-atom positions for each node are then generated using the predicted backbone frame, the backbone and side-chain torsion angles, as well as the predicted amino acid type (the amino acid type with the maximum probability). This calculation is done using the function *frames_and_literature_positions_to_atom14_pos* in the AF2 script *all_atom*.*py*.

The predicted all-atom positions are postprocessed to connect the unordered nodes into chains of amino acid sequences. This is done by using the heuristic algorithm in ModelAngelo(2). Intuitively, it connects nodes by minimizing the total length of peptide bond, based on the fact that the C-N distance in a peptide bond is less than 1.4 Å. Profile-sequence-based alignments between the predicted sequence profiles (i.e., *A* in Eq. 3) and the input sequences are further employed to correct the sequence assignment. Inspired by AF2, recycling is also used here by replacing the predicted Cα atoms in the first step by the ones from the postprocessed structure. Three rounds of recycling are used in this work.

#### The loss function for Cry-Net

The loss function here can be written as:

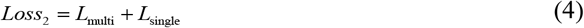

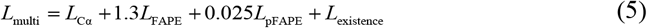

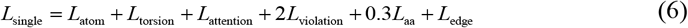

where the overall *Loss*_2_ can be divided into *L*_single_ (single-step loss) and *L*_multi_ (multi-step loss). The single-step loss is the final loss output after passing through 7 structure modules, while the multi-step loss is calculated 7 times within the 7 structure modules and averaged. They are described in detail below.

*L*_*atom*_ is the all-atom RMSD. It calculates the RMSD loss of all atoms on the main chain and side chain, which can be written as:

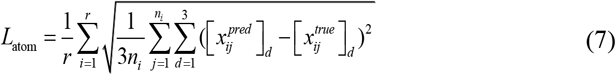

where *r* represents the total number of residues, *n*_*i*_ represents the number of atoms in the *i*-th residue, *x*_*ij*_ represents the atomic coordinates of the *j*-th atom in the *i*-th residue, and *d* represents the *d*-th dimension of the 3D coordinates.

*L*_torsion_ is the L_2_ distance between the torsion angles of the network output (i.e., *α* in Eq. 3) and the torsion angles of the deposited model. It is the same as the torsion angle loss defined in AF2.

*L*_attention_ is the classification loss when the nodes attend to the amino acids in the sequence. The calculation of this loss is the focal loss(57). The purpose of this loss is to ensure that each node can correctly match each amino acid in the sequence.

*L*_violation_ penalizes unreasonable bond angles and bond lengths in the model. It imposes certain constraints on the model’s output to ensure that the model’s output has ideal bond angles and lengths. The definition of this loss is the same as AF2.

*L*_aa_ represents the amino acid classification loss. This loss calculates the focal loss by comparing the twenty-dimensional amino acid probability vector (i.e., *A* in Eq. 3) and the one-hot vector corresponding to the amino acid identity in the deposited structure. The main purpose of this loss is to assign an amino acid identity to each node.

*L*_edge_ represents the edge classification loss. The edge classification is based on whether there is a connection between two nodes in the sequence. If there is a connection, we want them to be classified as neighbors. Specifically, this loss calculates the focal loss between *E* in Eq. 3 and the classification corresponding to the edge.

*L*_Cα_ is only the RMSD loss of Cα atoms. Its definition is similar to the all-atom RMSD (i.e., *n*_*i*_*=1* in Eq. 7).

*L*_FAPE_ comes from AF2, which plays a role in updating the backbone frame. It can be written as:

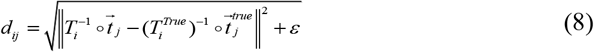

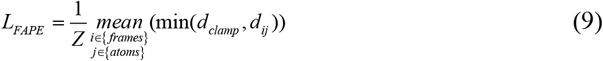

where *T*_*i*_ represents the *i*-th backbone frame (i.e., *T* in Eq. 3). *t*_*j*_ represents the coordinates of the *j*-th Cα atom. Both *d*_*clamp*_ and *Z* are taken as 10 Å here.

*L*_pFAPE_ is a simple regression loss. It calculates the mean squared error (MSE) loss between the predicted FAPE loss for each residue (i.e., *P* in Eq. 3) and the actual FAPE loss (i.e., Eq. 9). It is worth noting that in the post-processing, *P* will be normalized to a confidence score between 0 and 100 as the final confidence score.

*L*_existence_ represents the residue existence loss. Each residue can be classified into two classes: artificially added (see Supplementary Information S2.5) and naturally occurring. This loss calculates the focal loss between *M* in Eq. 3 and the corresponding one-hot vector of the residue class.

#### Software availability

The source code and model parameters are available at https://github.com/SBQ-1999/CryFold/ and https://yanglab.qd.sdu.edu.cn/CryFold/

## Acknowledgements

We would like to thank K. Jamali and S.H.W. Scheres for their kind help in the benchmark of ModelAngelo; S. Caffarri, A. Naschberger, R. Sobotka, and F. Wu for help with the structural studies. This work is supported by the Ministry of Science and Technology of the People’s Republic of China (2024YFA0916901), the National Natural Science Foundation of China (NSFC T2225007, T2222012, 32430063), the Fundamental Research Funds for the Central Universities, European Research Council (ERC-805230), and Westlake Visiting Professor Fellowship to AA.

## Supplementary Information

### S1. Evaluation metrics

We used the same set of evaluation metrics defined by ModelAngelo(1), including *backbone recall, backbone precision, backbone RMSD, Cα RMSD, amino acid accuracy*, and *completeness*.

- *Backbone recall* is the fraction of deposited residues (represented by Cα atoms) that have a predicted residue (represented by Cα atom) within 3 Å.
- *Backbone precision* is the fraction of predicted residues (represented by Cα atoms) that have a deposited residue (represented by Cα atom) within 3 Å.

The remaining subsequent metrics involve the matching between deposited residues and predicted residues, and only consider the deposited residues that have a predicted residue within 3 Å. We first calculate the distances between the Cα atoms in predicted structure and the Cα atoms in deposited structure. The Hungarian Matching Algorithm(2) is then used to match the predicted residues with the deposited residues such that the total distance is minimized. The metrics can be defined based on the matched residue pairs.

Let *p* represent the number of matched residues that have the same amino acid identity; *m* represent the total number of matched residues; and *n* represent the total number of residues in the deposited structure.

- *Cα RMSD* is the root-mean-square deviation (RMSD) between the Cα atoms of the matched residue pairs.
- *Backbone RMSD* is similar to the former but includes all four main-chain atoms (Cα, C, O, and N).
- *Amino acid accuracy* is the fraction of residue pairs that share identical amino acid types (i.e., *p/m*).
- *Completeness* is the fraction of all deposited residues (including those unmatched residues) that have a matched predicted residue with the same amino acid identity (i.e., *p/n*). It is worth noting that *Completeness ≈ Backbone recall × Amino acid accuracy*.

### S2. Network details

#### S2.1 U-Net

The encoder of the U-Net (3) architecture adopts a bottleneck structure (4), which is the same as the convolutional neural network (CNN) module of Cry-Net. The decoder uses the Res2Net architecture (5, 6), as shown in Fig. S12. The use of bottleneck for downsampling aims to preserve as much spatial information as possible, while the use of Res2net for upsampling aims to extract as much semantic information as possible. The spatial information and semantic information are then combined to predict the probability map of Cα atoms.

#### S2.2 U-Net training

As mentioned in S2.1, the encoder architecture of U-Net is the same as the CNN network structure in Cry-Net. In this case, we employed a form of transfer learning technique, where we used the pre-trained CNN network parameters from the Cry-Net as the initial weights for the encoder of U-Net, without increasing the training sample size. This is because the Cry-Net was thoroughly trained on various protein-related knowledge, such as structure prediction, amino acid classification, and confidence prediction. Therefore, the CNN network from Cry-Net can extract richer semantic information from the cryo-EM density maps, which cannot be solely obtained through the task of predicting Cα positions in Stage 1. The encoder architecture of U-Net can extract richer information through this pre-training process.

#### S2.3 Sequence attention and IPA

The Sequence Attention and IPA modules are only briefly mentioned in the main text. In this section, they will be presented in the form of pseudocodes.

##### Algorithm Si

Sequence attention

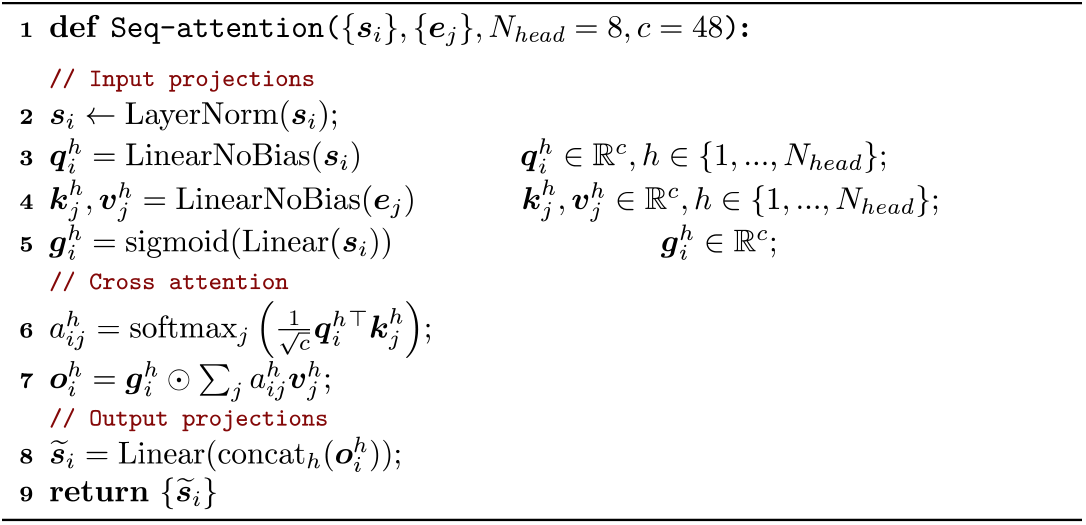

In this algorithm, the input ***s***_*i*_ represents the node representation of the *i*-th node, and the input ***e***_*j*_ represents the ESM-2 (7) embedding representation of the *j*-th amino acid in the sequence. The output 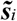 is the updated node representation of the *i*-th node.

##### Algorithm S2

Local invariant point attention with 3D-RoPE

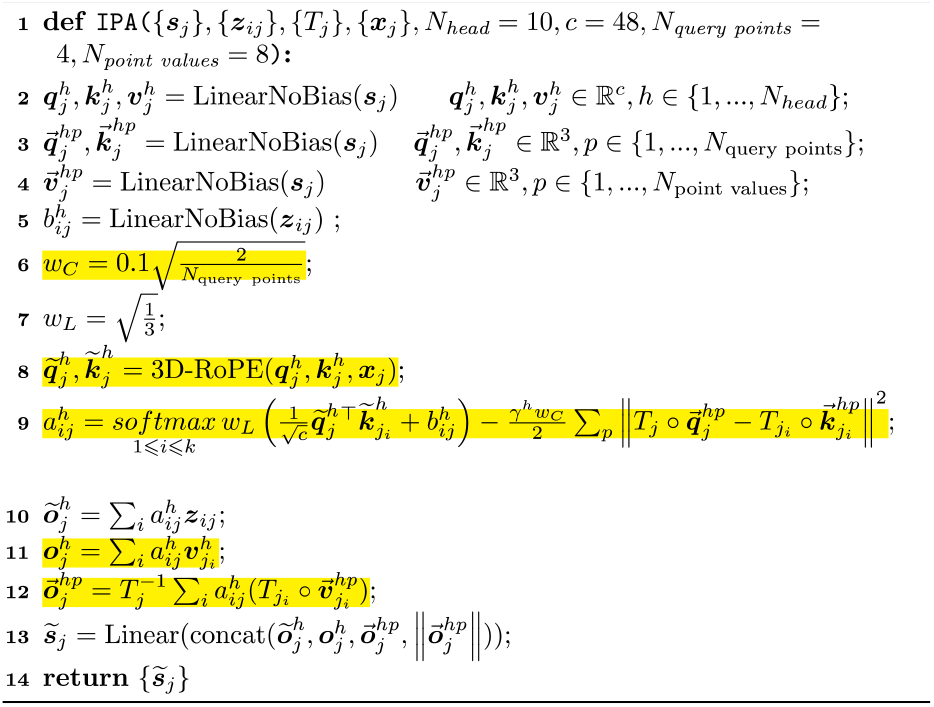

In this algorithm, the input ***s***_*j*_ represents the node representation of the *j*-th node, ***z***_*ij*_ represents the edge representation between the *j*-th node and its *i*-th neighbor, *T*_*j*_ represents the backbone frame of the *j*-th node, and ***x***_*j*_ represents the position (i.e., the coordinates of the Cα atom) of the *j*-th node. The output 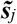 is the updated node representation of the *j*-th node. The highlighted text here outlines the main differences between the IPA module in this work and the one in AF2 (8). The attention used here is a local form and includes a 3D rotary position embedding (see below).

#### S2.4 3D Rotary Position Embedding (3D-RoPE)

3D-RoPE is used in the node attention and IPA modules of the Cry-Net. To describe this in detail, let’s first introduce the 1D rotary position encoding (9):

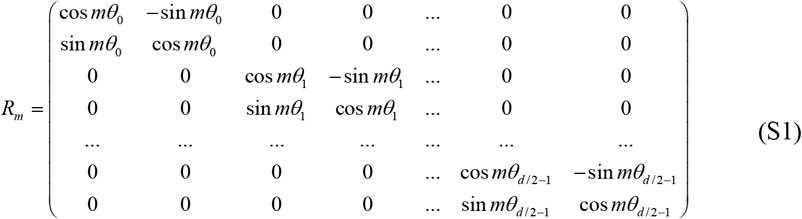

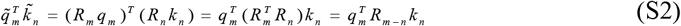

where *q* and *k* represent the query vector and the key vector used in the attention mechanism, respectively. They are both *d*-dimensional vectors. The above formula is to introduce the positions *m* and *n* of the sequence into the attention mechanism using the multiplicative position encoding. We extend this process to 3D space:

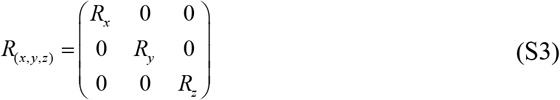

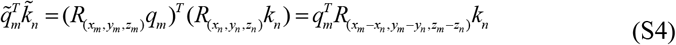

Here, we assume that the dimensions of the query and key vectors are both multiples of three, and (*x, y, z*) represents the position of each node (i.e., the coordinates of the Cα atoms). The 3D position encoding has richer connotations compared to the 1D position encoding, as it not only considers the distance information between nodes but also the directional information between nodes. Furthermore, this attention mechanism can actually be applied along the three coordinate axes:

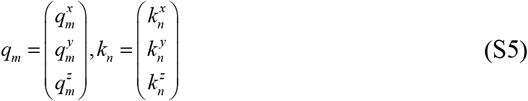

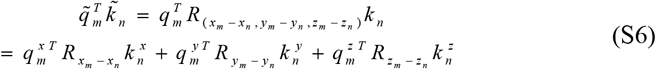

The first one-third of the dimensions in the query and key vectors represent the information along the *x*-axis, the middle one-third represent the information along the *y*-axis, and the last one-third represent the information along the *z*-axis. From this perspective, the 3D-RoPE attention can be seen as the sum of three 1D-RoPE attentions in different directions (*x, y, z*). The 3D-RoPE attention is realized by the inner product calculation. In the 3D world, the inner product has a clear physical meaning, such as representing the work done:

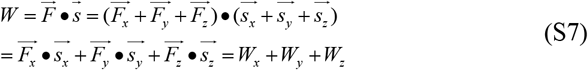

The total work *W* is the inner product of the force 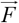 and the displacement 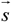. From another perspective, the total work *W* can be decomposed into the work done along the three coordinate axes. This has an analogy with the 3D-RoPE attention.

If we interpret the 3D-RoPE attention as the interaction between residues, then based on the long-term decay property of the 1D-RoPE(9), the 3D-RoPE, which can be seen as the sum of three 1D-RoPE, will also exhibit long-range decay. In this way, the 3D attention between residues will weaken as the distance increases, which nicely captures the mechanism of the inter-residue interactions.

Finally, 3D-RoPE is applied together with local attention, which ensures good extrapolation capabilities. Specifically, the number of tokens processed in each attention operation during both the training and testing phases remains consistent (i.e., limited to the nearest *k* residues in space). Even when the spatial dimensions of a tested map become very large, the local attention constrains each residue to attend to its neighboring residues only. As a result, the relative positional distances in 3D-RoPE are in a limited range, even for large maps.

#### S2.5 Cry-Net training

For the convenience of narration, we restate the Eq. 3 from the main text here. The input and output of Cry-Net in each the *n*-th iteration are defined by the following equation:

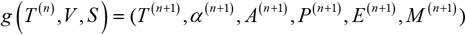

The frame *T* can be defined as (*R, t*), where *R*∈ ℝ^*r*×3×3^ and *t* ∈ ℝ^*r*×3×1^. At the beginning of the training, *t*^(0)^ is initialized as the Cα atomic coordinates of the training data after adding Gaussian noise 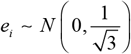 along each dimension, and the rotation matrix *R* ^(0)^ is randomly initialized (1).

During the training process, a Cα atom is randomly selected from each PDB (10) data sample, and the 200 nearest residues in the spatial region are cropped. Inspired by ModelAngelo, 10% of the residues are randomly replaced with peptide chains of length 2-5. The purpose of this is to simulate the potential redundancy in the Cα atom coordinates output by U-Net, i.e., there may be no corresponding residue with deposited structure around the output nodes. In this case, the output *M* of the network is used to determine whether the nodes output by U-Net are redundant or not.

#### S3. Supplementary figures

**Fig. S1.**
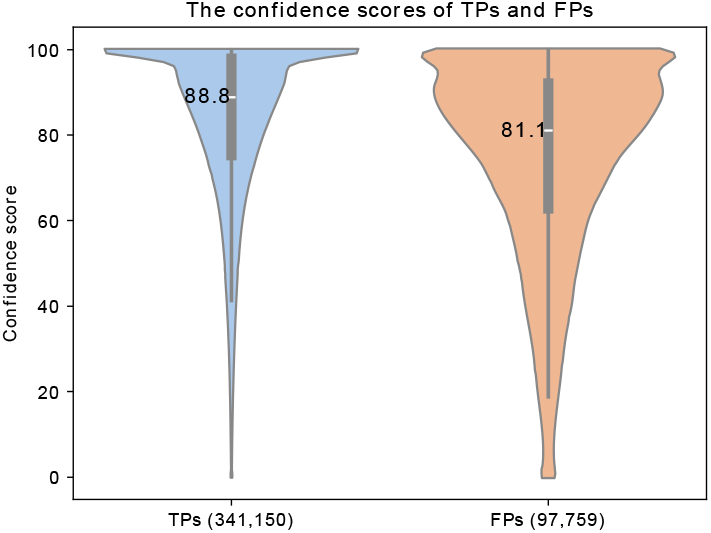
The distribution of confidence scores of the CryFold models for the 177 maps. TPs: true positives, FPs: false positives. Note that the FPs are possible to be misjudgments, as illustrated by the example in Fig. 3.

**Fig. S2.**
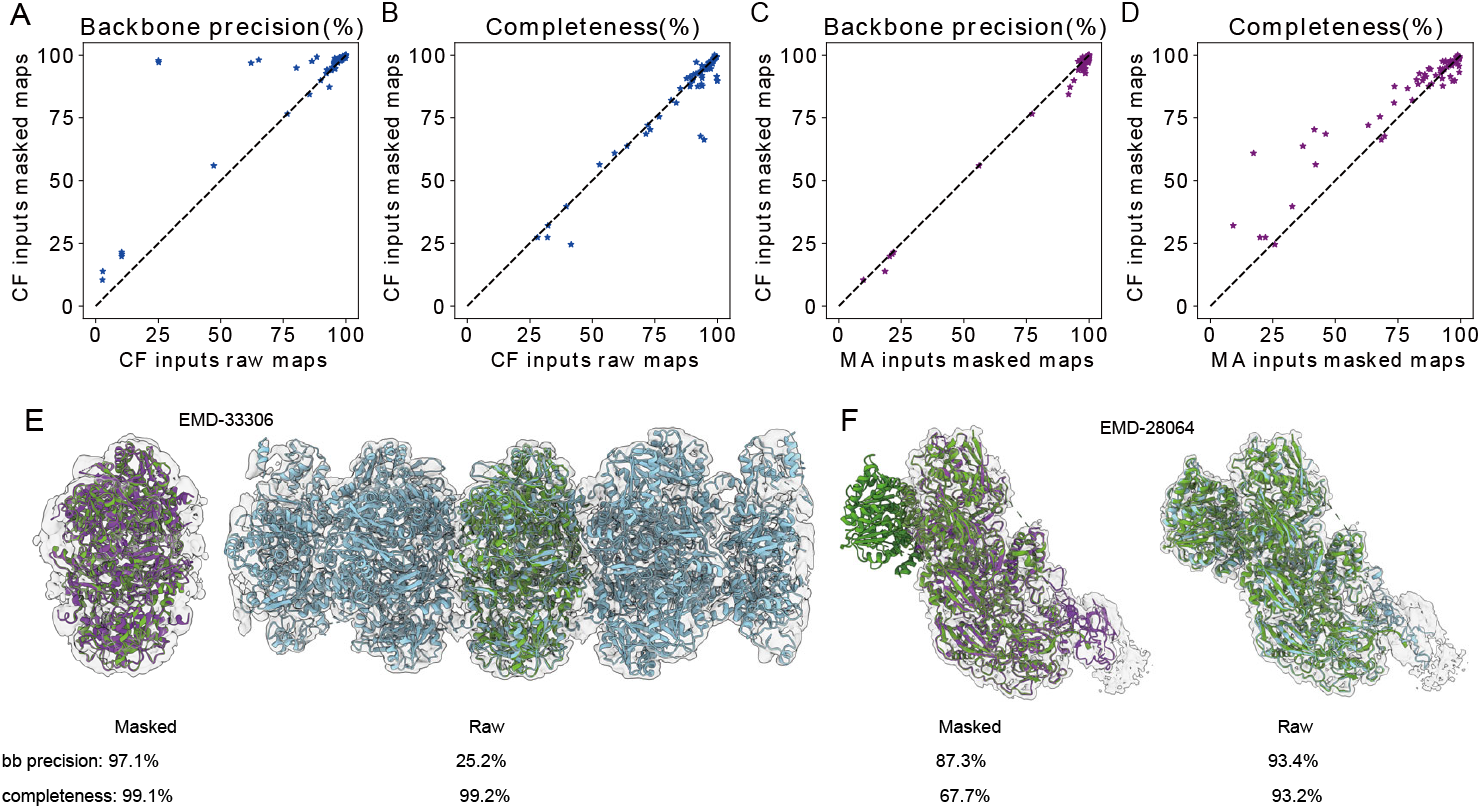
The impact of map masking. (A) Head-to-head comparison of the model backbone precision. (B) Head-to-head comparison of the model completeness. (C)/(D) Head-to-head comparison of the model backbone precision/completeness between CryFold and ModelAngelo by inputting masked maps. (E) Appropriate masking can screen out regions unrelated to the deposited structure protein. The gray surface is the density map. The green cartoon represents the deposited structure. The purple/blue cartoon is the CryFold model constructed using the masked/raw map. (F) Inappropriate masking can lead to incorrect atomic model. The coloring scheme is the same as in (E).

**Fig. S3.**
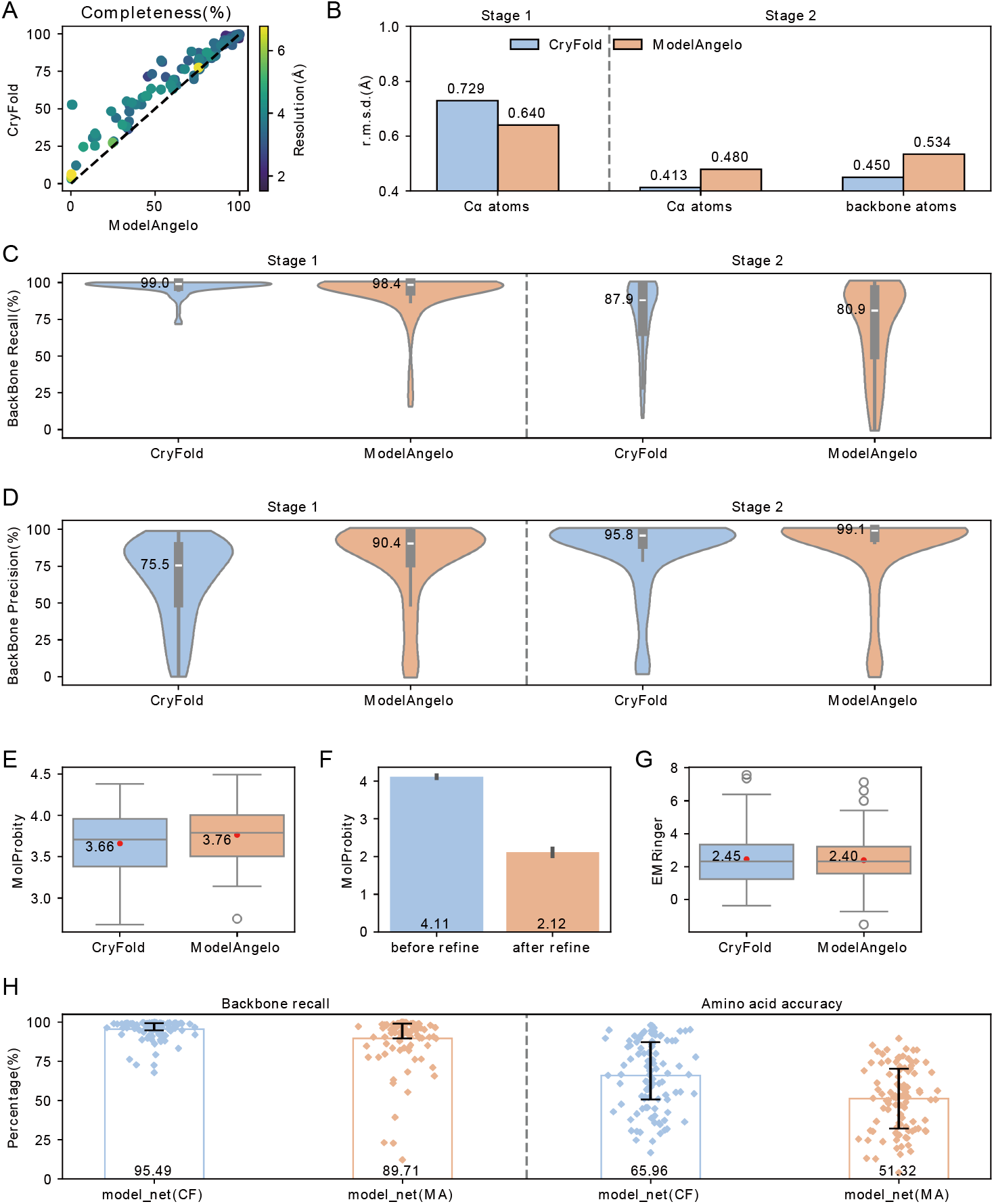
Comparison of CryFold and ModelAngelo on 101 non-redundant density maps. (A) Head-to-head comparison of the model completeness. (B) Bar plots comparing the average RMSDs of Cα atoms and backbone atoms. Stage 1 predicts only the Cα atoms, while Stage 2 outputs all-atom models. (C)/(D) Violin plots of the backbone recall/precision. (E) Box plots of the MolProbity score, in which the red dots represent the mean values. (F) Results of the 21 CryFold predicted models with MolProbity scores > 4 before and after relaxation. (G) Box plots of EMRinger, in which the red dots represent the mean values. (H) Backbone recall and amino acid accuracy of the intermediate models.

**Fig. S4.**
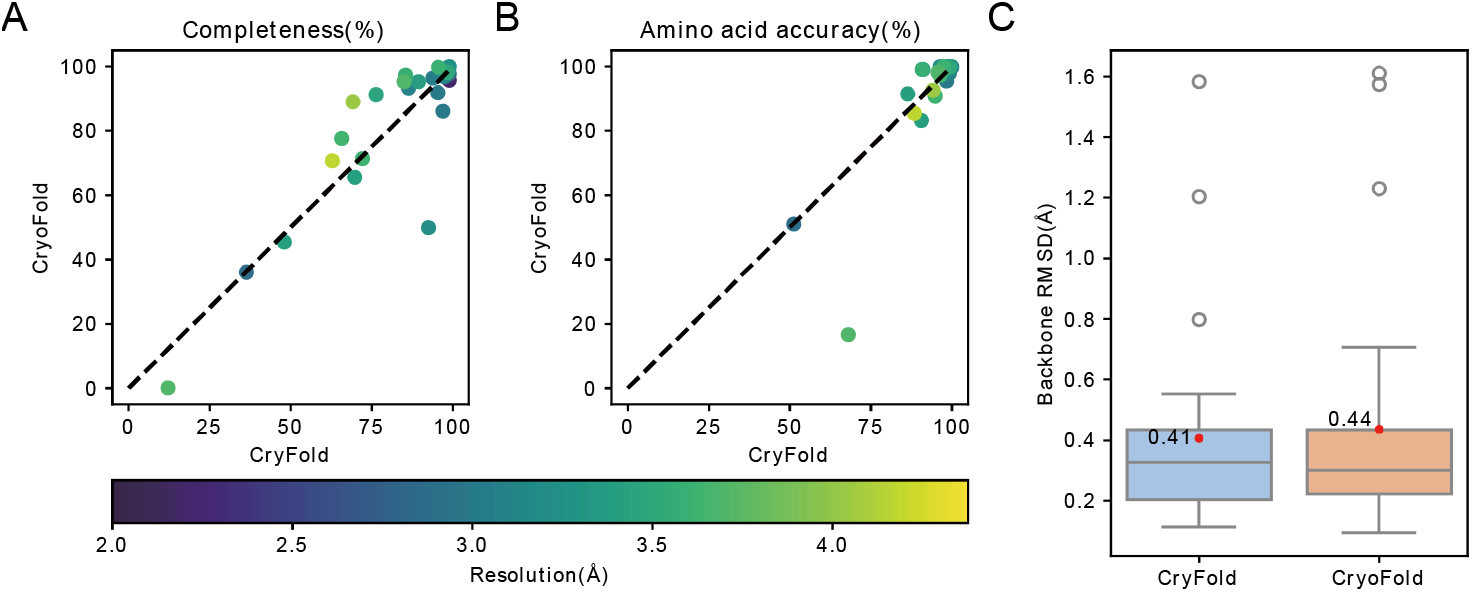
Comparison of CryFold and CryoFold (https://cryonet.ai/cryofold) on 25 randomly selected maps from the non-redundant test set. (A)/(B) Head-to-head comparison of the model completeness/amino acid accuracy. (C) Box plots comparing the backbone RMSD. The red dots represent the mean.

**Fig. S5.**
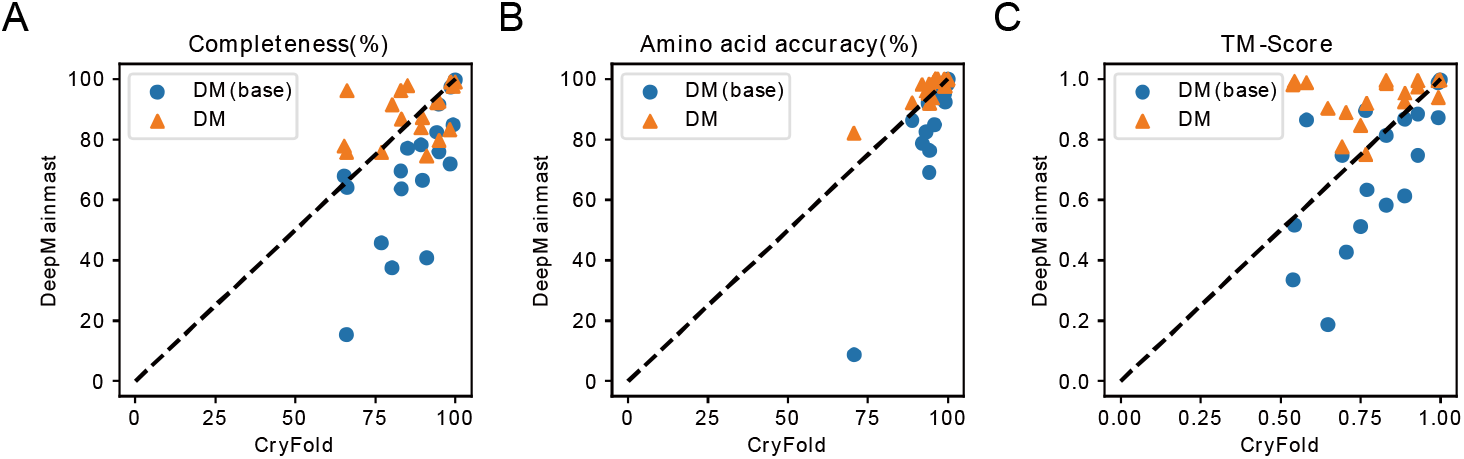
Comparison of CryFold and DeepMainmast (11) on 17 test density maps. The maps and results are cited from the DeepMainmast paper. (A-C) Head-to-head comparison of the model completeness, amino acid accuracy and TM-Score. DM(base) refers to running DeepMainmast without using AlphaFold. DM refers to running DeepMainmast with AlphaFold.

**Fig. S6.**
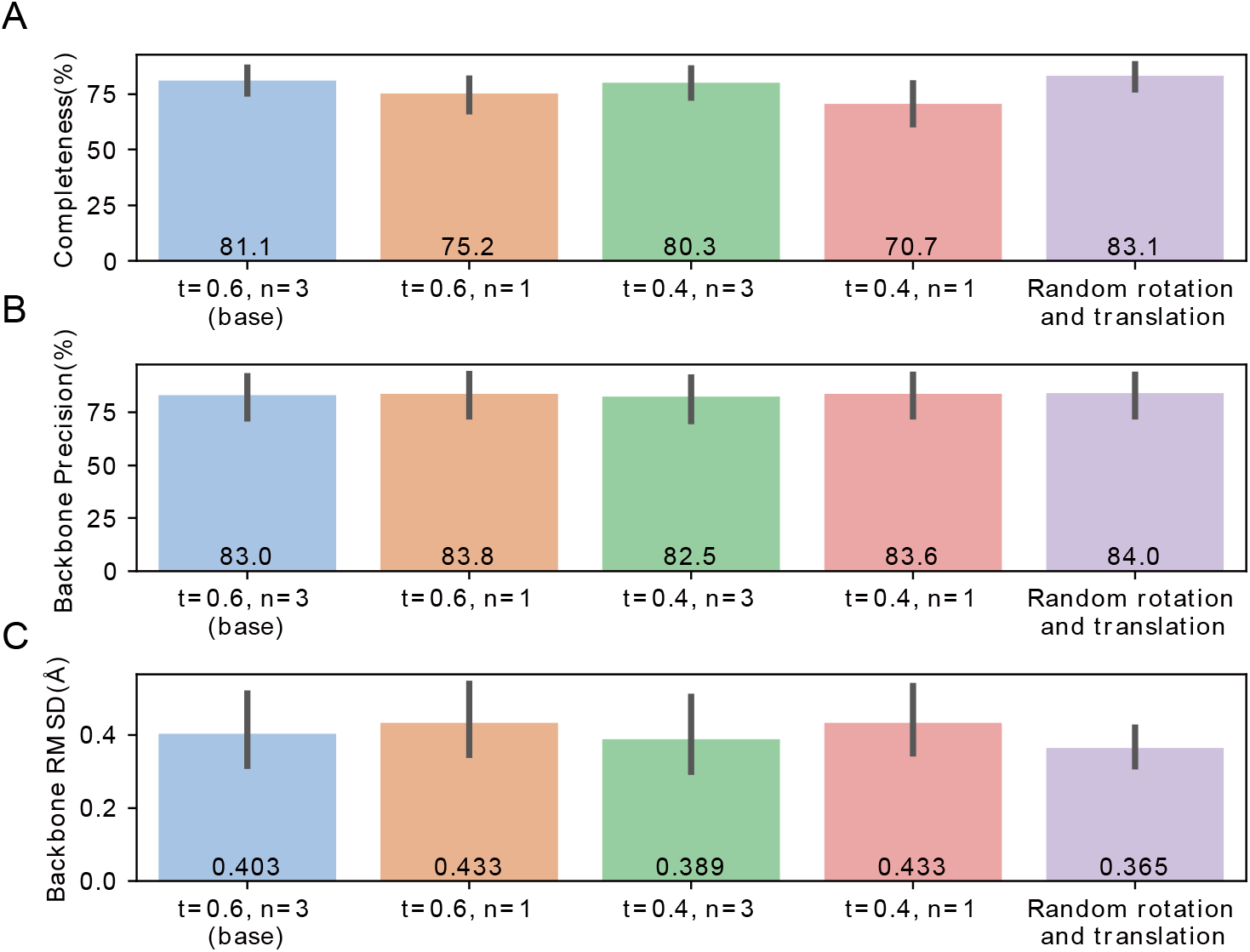
Randomly selected 33 density maps from the non-redundant test set to test CryFold’s sensitivity to hyperparameters and rotation-translation. The parameter *t* represents the threshold for predictions made by the classification network in Stage 1. The parameter *n* represents the number of recycling rounds in Stage 2. Random rotation and translation refer to the density maps. CryFold will use the transformed density map as input and compare the results with the native model. (A-C) Bar plots comparing the average completeness, backbone precision, backbone RMSD.

**Fig. S7.**
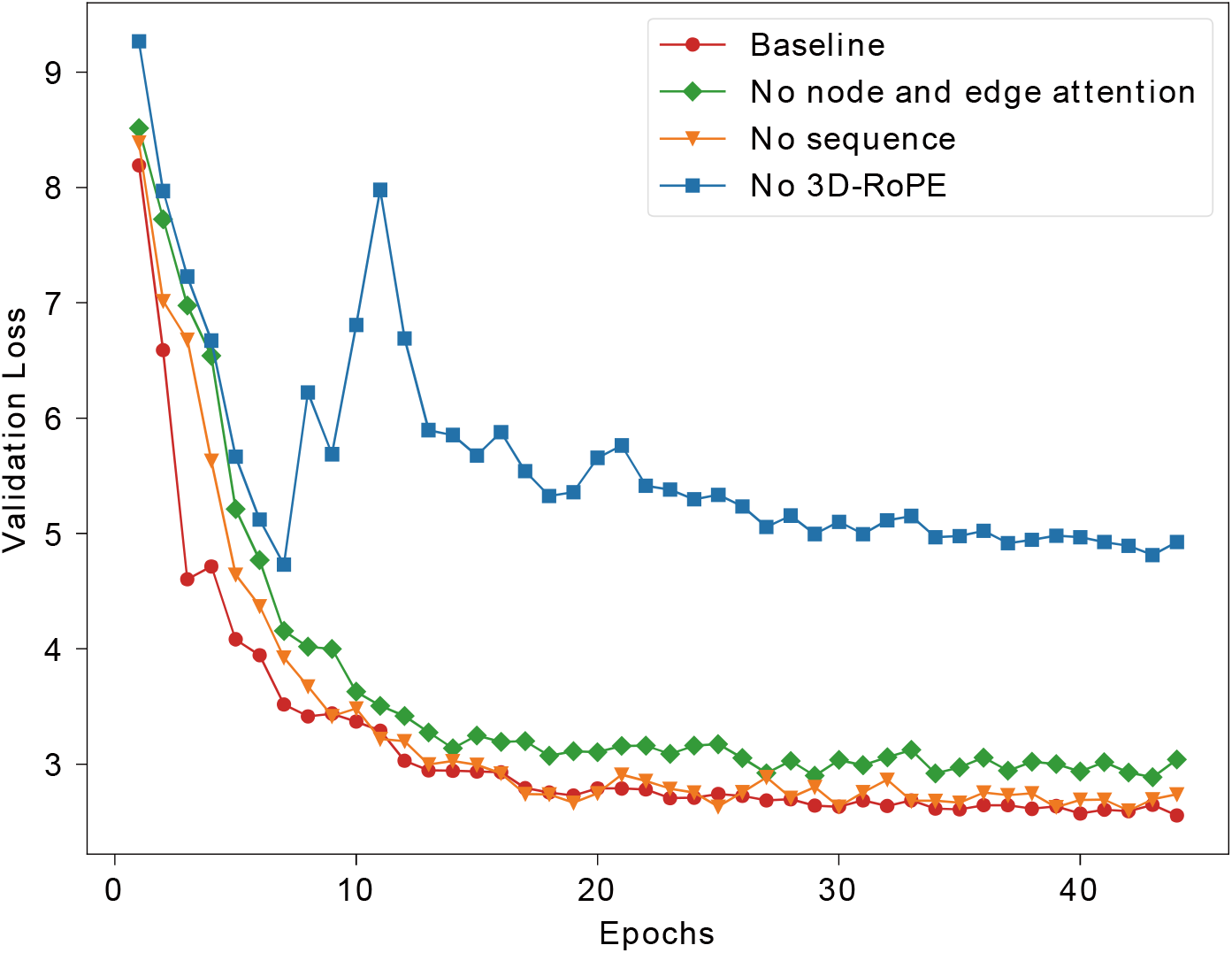
The curves of loss function under four different configurations.

**Fig. S8.**
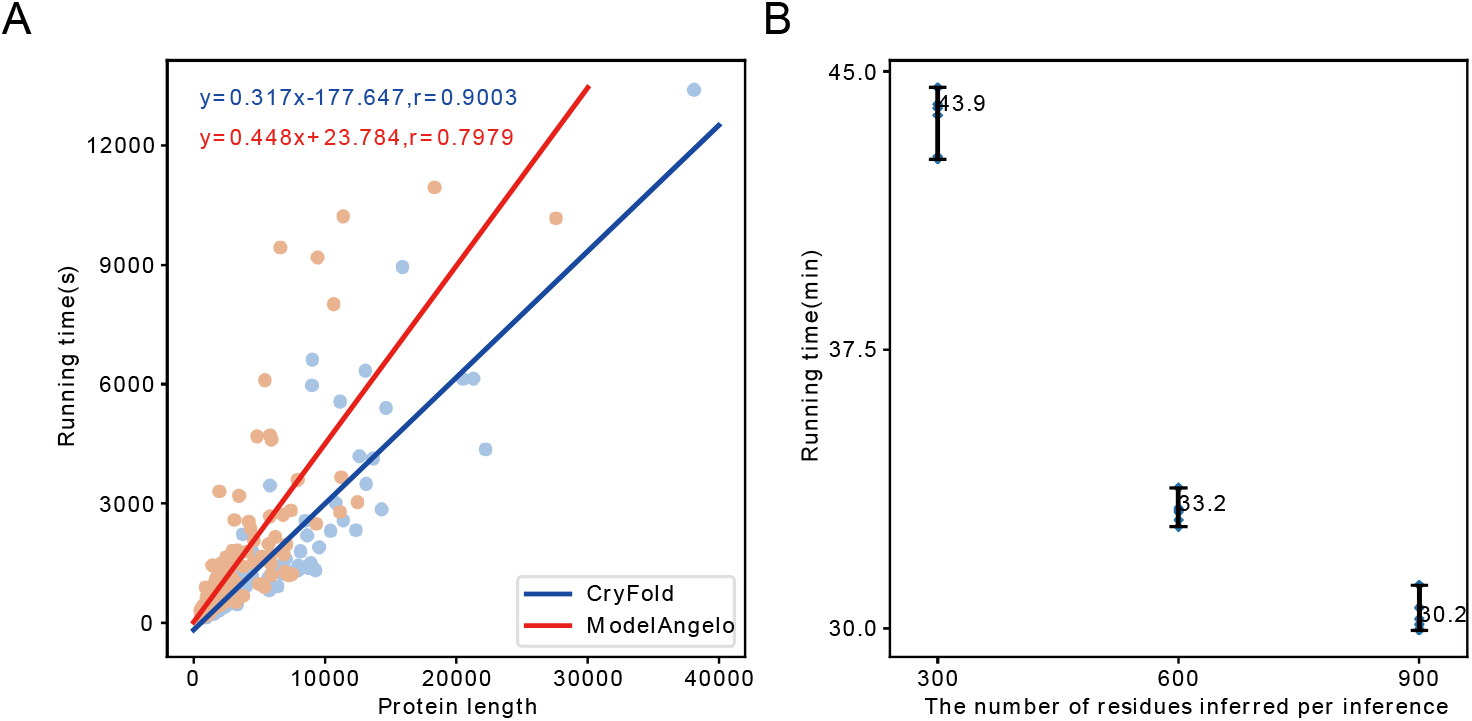
Computational efficiency of CryFold. (A) Running time for building 177 maps using CryFold and ModelAngelo. (B) Running time for an example protein (EMD-29327, PDB ID: 8FNV) under different memory configurations (15GB, 19GB, 24GB). To eliminate the effects of randomness, each configuration was run six times.

**Fig. S9.**
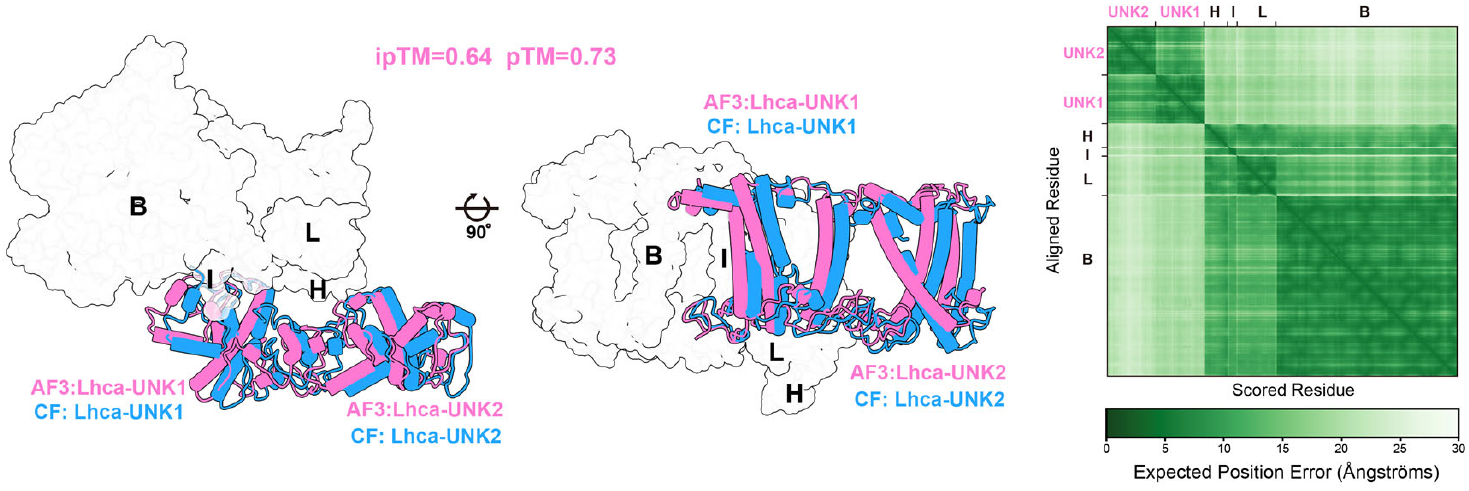
Modelling evaluation by the CryFold and AlphaFold3 of newly identified proteins (Related to Fig. 6B, 6C). Superposition of the CryFold and AlphaFold3 models of a subcomplex comprising chains B, H, I, L, newly identified Lhca-UNK1, Lhca-UNK1. While the core subunits superimpose well (gray surface), the positions of the newly modeled proteins are shifted (cartoon). The figure on the right-hand side is the Predicted Aligned Error (PAE) plot of the model predicted by AF3 (12).

**Fig. S10.**
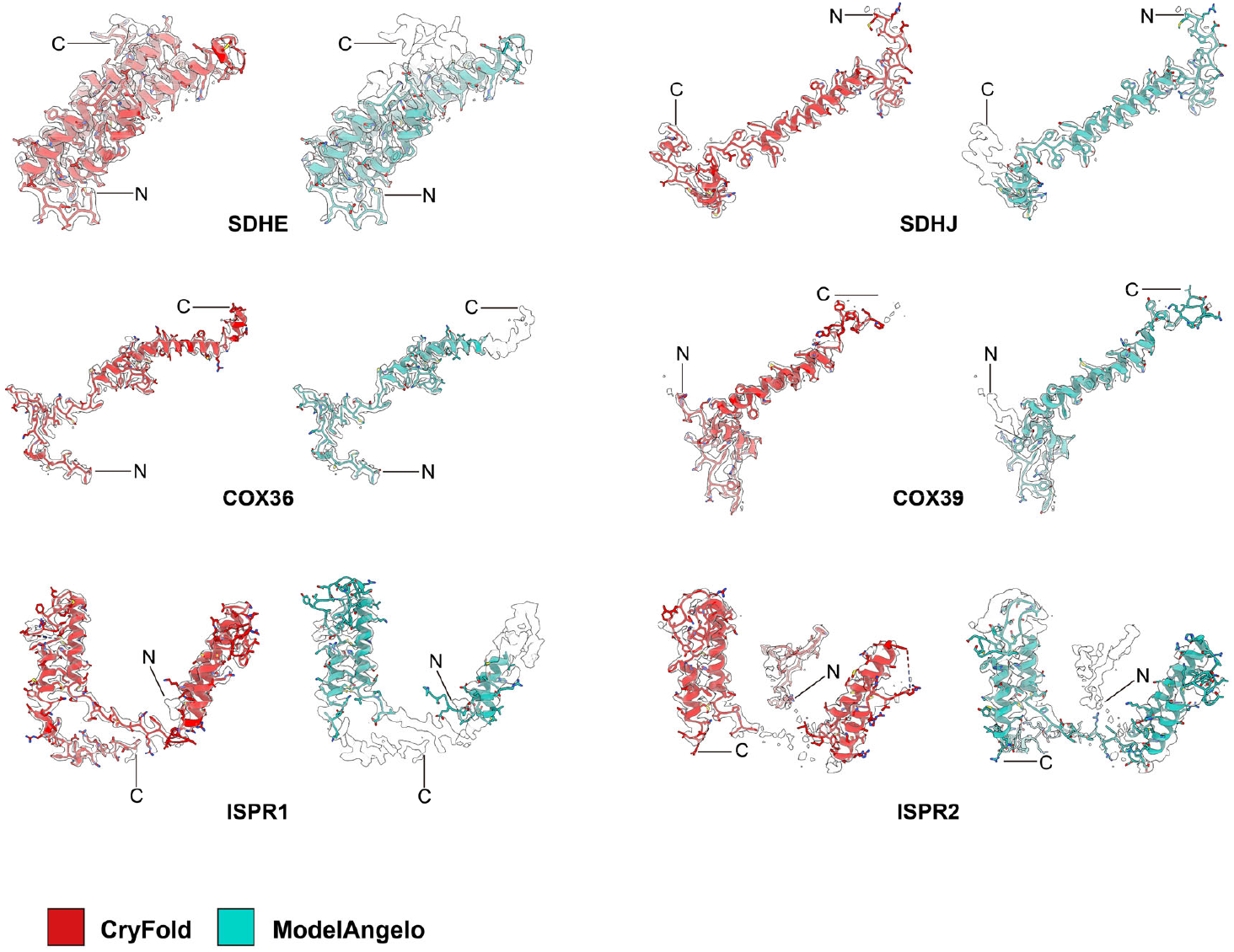
Modelling evaluation by the CryFold and ModelAngelo in low-resolution regions. Six protein models of the mitochondrial respirasome II2-III_2_-IV2 (13) are compared and shown with their corresponding densities. Termini are indicated for each protein.

**Fig. S11.**
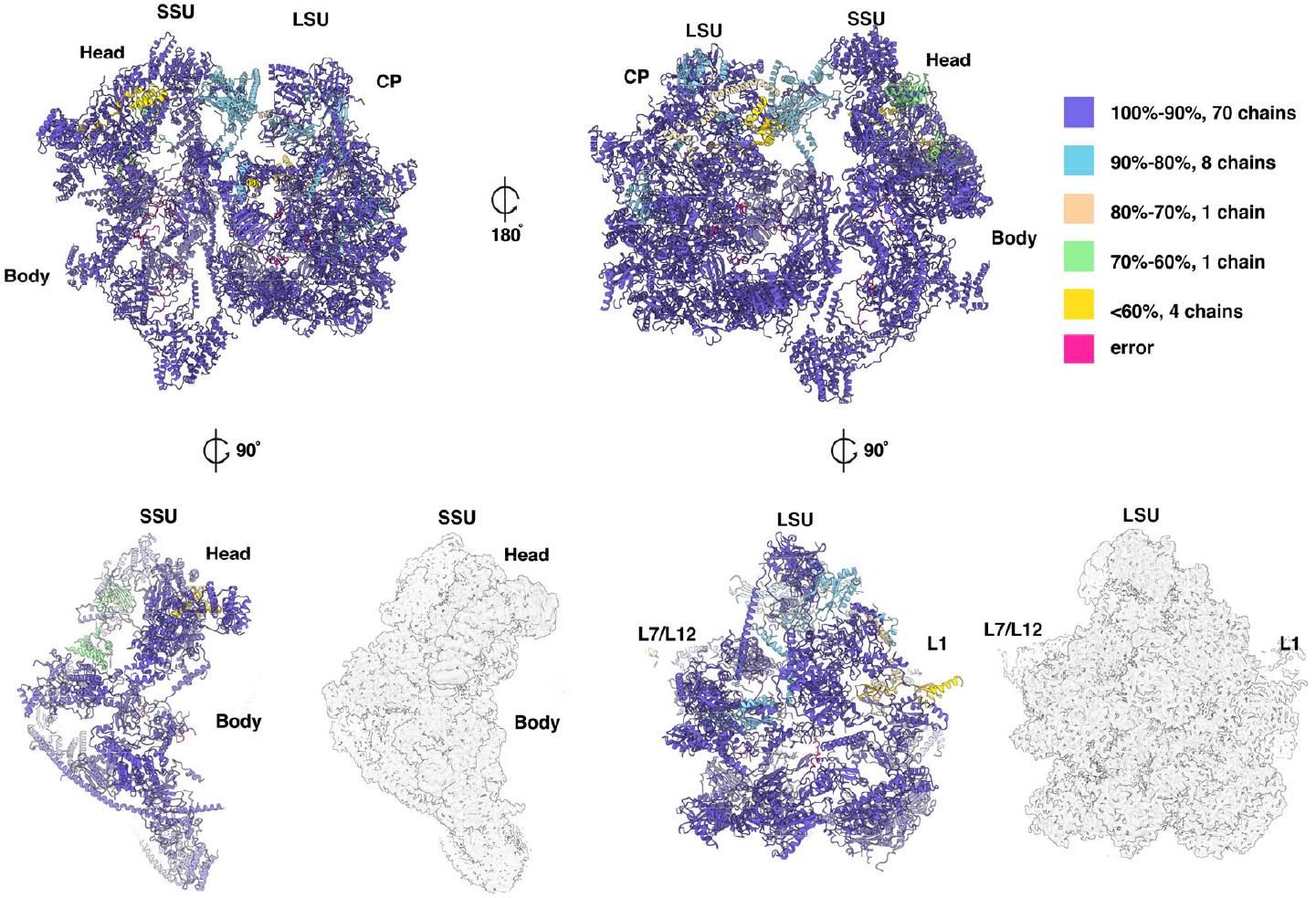
Modelling of the mitoribosome. Overall view of the CryFold model of the human mitoribosome, and its individual subunits. Proteins are colored according to model completeness. The corresponding density map (EMD-13980) is shown for individual subunits.

**Fig. S12.**
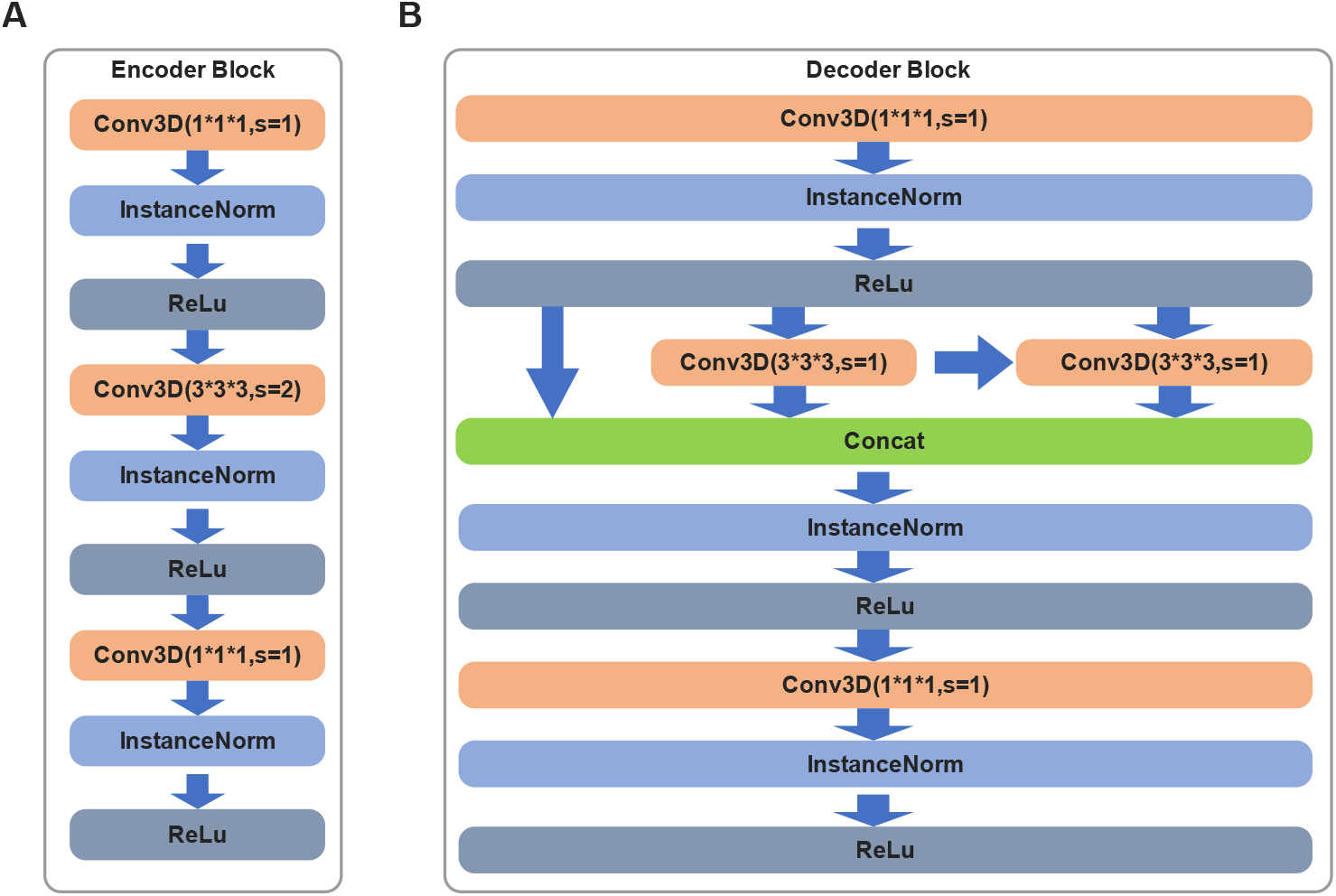
Architecture of U-net. (A) Encoder module and the downsampling module are integrated together, and a bottleneck architecture(4) is adopted. (B) Decoder module adopts the Res2Net architecture (5, 6).

**Fig. S13.**
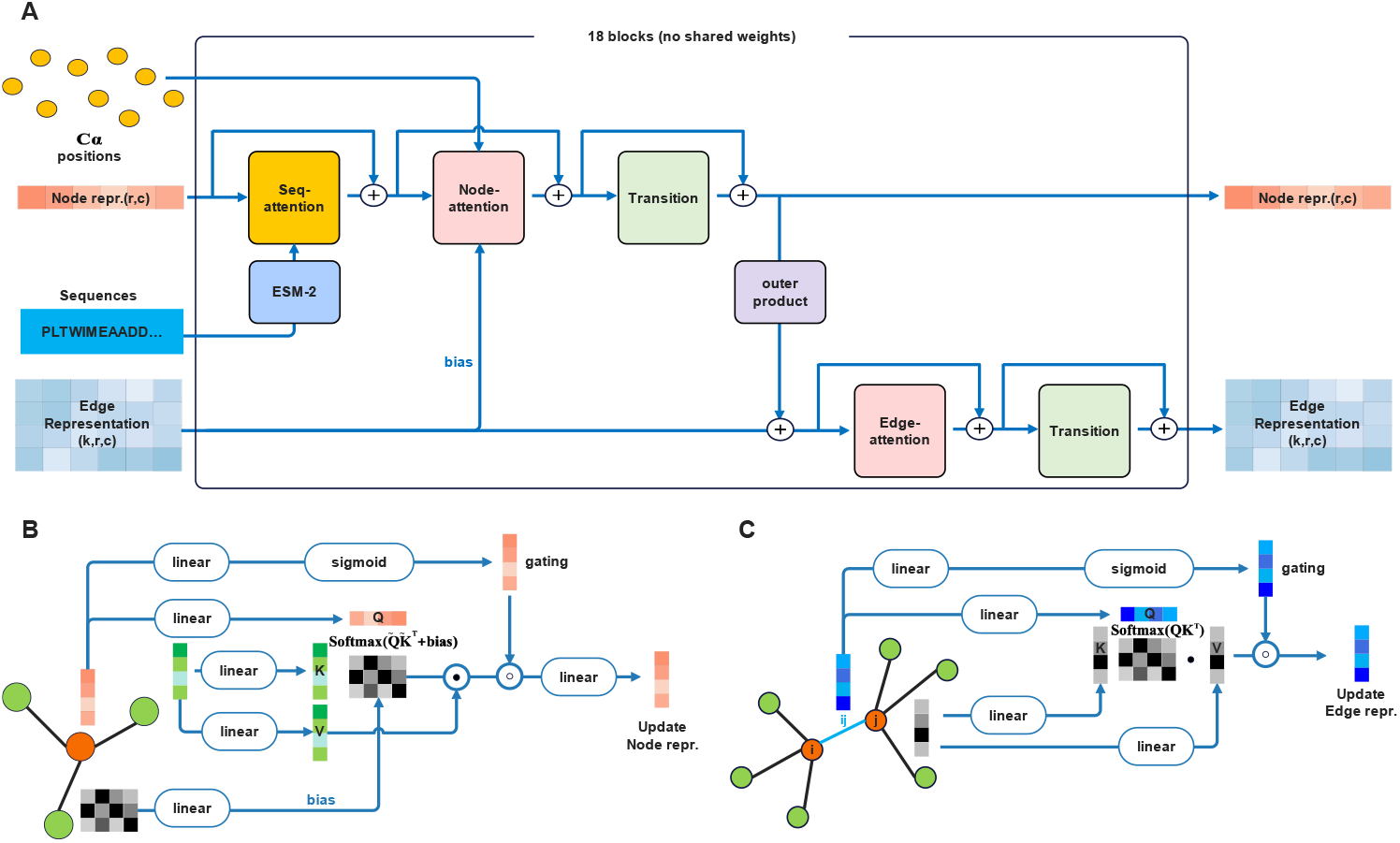
Architectural details of the encoder network. (A) Cryformer module. (B) node attention layer. (C) edge attention layer.

**Fig. S14.**
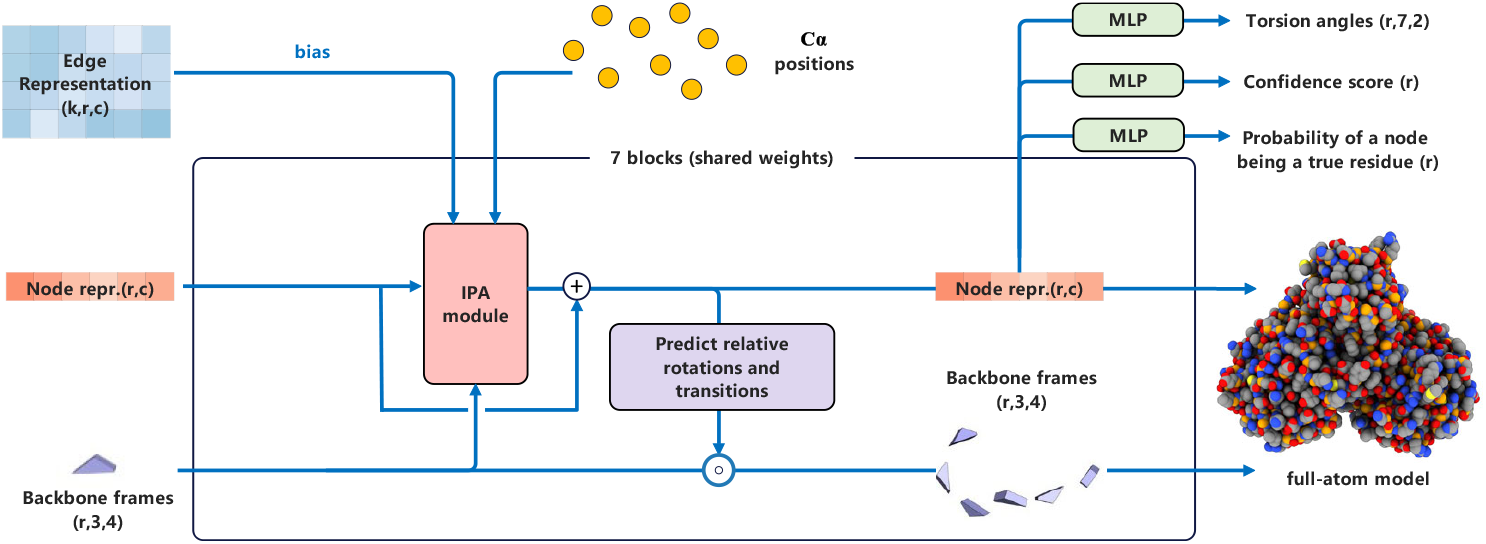
Architectural details of the decoder network Structure Module. This module is similar to the structure module in AF2 (8). The difference is that the input here includes additional node position information (i.e., the Cα positions). The updated node representations are passed through separate MLPs to output *T, α, P, M* in Eq. 3.

**Fig. S15.**
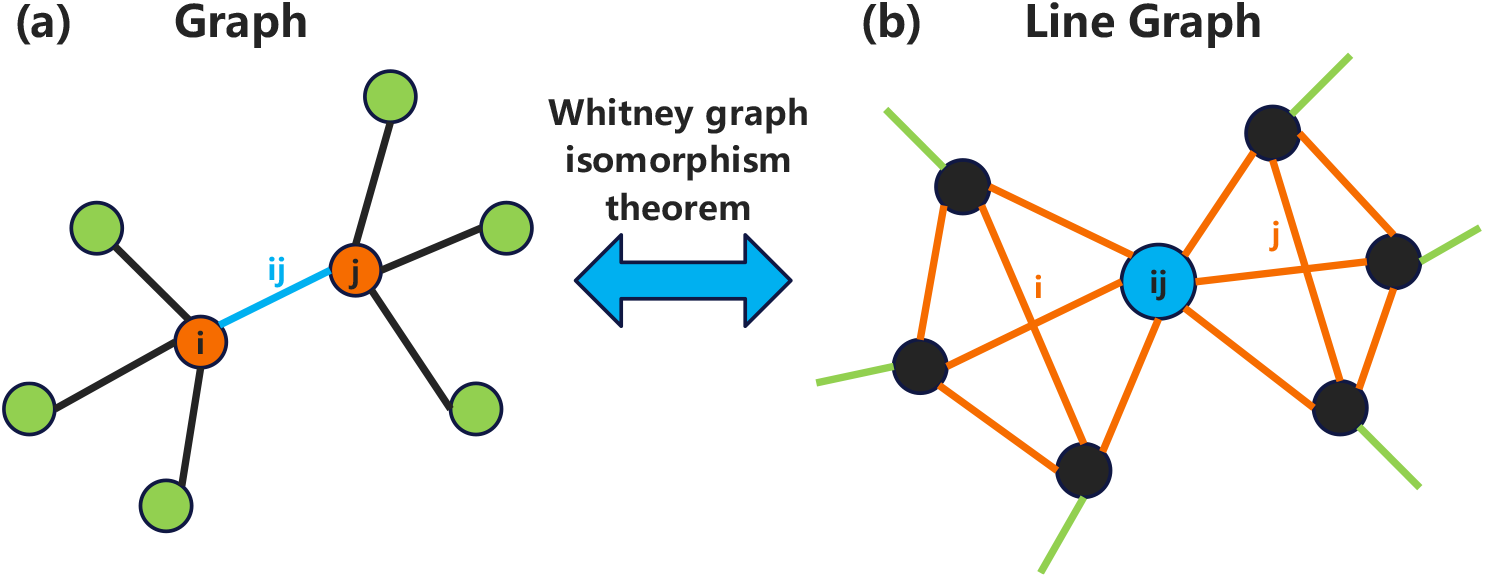
Relationship between node attention and edge attention. A graph (a) can be converted into a line graph (b), and vice versa. According to the Whitney graph isomorphism theorem, except for a very small number of special cases, the line graph uniquely determines the original graph. These two types of graphs have a certain duality. The edge attention to the edge *ij* of the graph on the left is equivalent to the node-attention to the *ij* node of the line graph on the right.

**Fig. S16.**
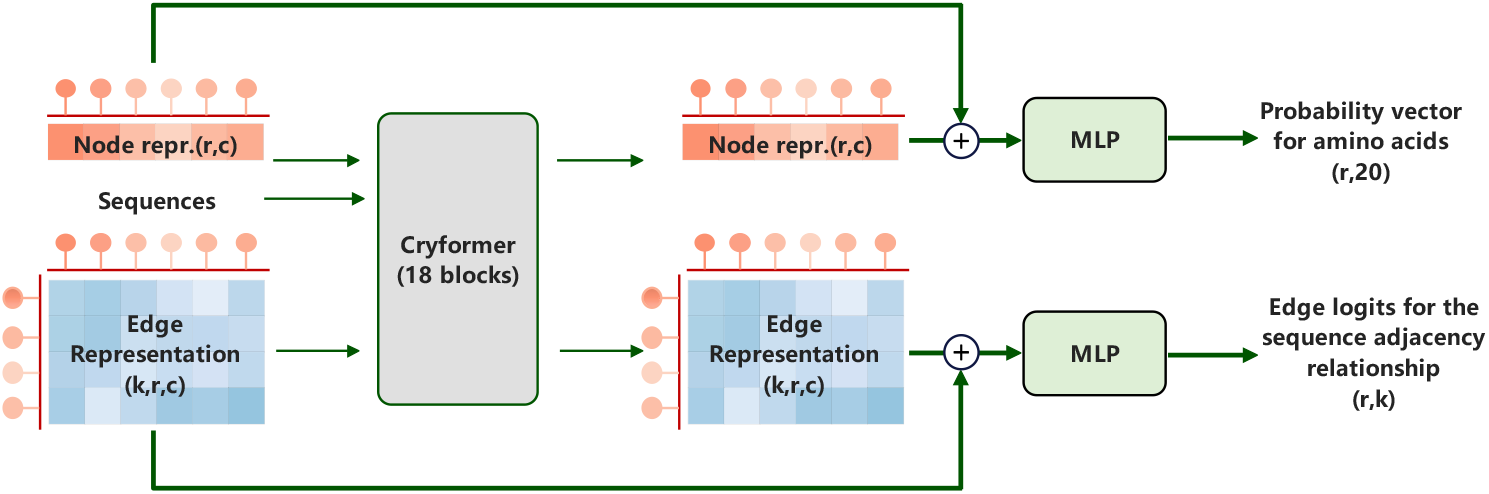
Detailed information for the Cryformer Module. The original node representation and the updated node representation are concatenated and passed through a separate MLP to predict the probability vector over the 20 types of amino acids for each node (i.e., *A* in Eq. 3). The original edge representation and the updated edge representation are concatenated and passed through another separate MLP to predict the edge connectivity (i.e., *E* in Eq. 3).

## References

1. Jumper J, et al. (2021) Highly accurate protein structure prediction with AlphaFold. Nature 596(7873):583–589.

2. Jamali K, et al. (2024) Automated model building and protein identification in cryo-EM maps. Nature 628(8007):450–457.

3. Nakane T, et al. (2020) Single-particle cryo-EM at atomic resolution. Nature 587(7832):152–156.

4. Amunts A, et al. (2014) Structure of the yeast mitochondrial large ribosomal subunit. Science 343(6178):1485–1489.

5. Kühlbrandt W (2014) Biochemistry. The resolution revolution. Science 343(6178):1443–1444.

6. Burley SK, et al. (2023) RCSB Protein Data Bank (RCSB.org): delivery of experimentally-determined PDB structures alongside one million computed structure models of proteins from artificial intelligence/machine learning. Nucleic Acids Research 51(D1):D488–D508.

7. Chua EYD, et al. (2022) Better, Faster, Cheaper: Recent Advances in Cryo-Electron Microscopy. Annu Rev Biochem 91:1–32.

8. de la Rosa-Trevin JM, et al. (2024) EMhub: a web platform for data management and on-the-fly processing in scientific facilities. Acta Crystallogr D Struct Biol.

9. Stuart DI, Subramaniam S, & Abrescia NG (2016) The democratization of cryo-EM. Nat Methods 13(8):607–608.

10. Yang J, et al. (2015) The I-TASSER Suite: protein structure and function prediction. Nat Methods 12(1):7–8.

11. Emsley P, Lohkamp B, Scott WG, & Cowtan K (2010) Features and development of Coot. Acta Crystallogr D Biol Crystallogr 66(Pt 4):486–501.

12. Croll TI (2018) ISOLDE: a physically realistic environment for model building into low-resolution electron-density maps. Acta Crystallogr D Struct Biol 74(Pt 6):519–530.

13. Brown A, et al. (2017) Structures of the human mitochondrial ribosome in native states of assembly. Nat Struct Mol Biol 24(10):866–869.

14. Muhleip A, et al. (2021) ATP synthase hexamer assemblies shape cristae of Toxoplasma mitochondria. Nat Commun 12(1):120.

15. Muhleip A, et al. (2023) Structural basis of mitochondrial membrane bending by the I-II-III(2)-IV(2) supercomplex. Nature 615(7954):934–938.

16. Wú F, et al. (2024) Structure of the II2-III2-IV2 mitochondrial supercomplex from the parasite Perkinsus marinus. bioRxiv:2024.2005.2025.595893.

17. Brown A, et al. (2015) Tools for macromolecular model building and refinement into electron cryo-microscopy reconstructions. Acta Crystallogr D Biol Crystallogr 71(Pt 1):136–153.

18. Chojnowski G, et al. (2022) findMySequence: a neural-network-based approach for identification of unknown proteins in X-ray crystallography and cryo-EM. IUCrJ 9(Pt 1):86–97.

19. Wlodawer A, Li M, & Dauter Z (2017) High-Resolution Cryo-EM Maps and Models: A Crystallographer’s Perspective. Structure 25(10):1589–1597 e1581.

20. Gao Y, Thorn V, & Thorn A (2023) Errors in structural biology are not the exception. Acta Crystallogr D Struct Biol 79(Pt 3):206–211.

21. Chen M, Baldwin PR, Ludtke SJ, & Baker ML (2016) De Novo modeling in cryo-EM density maps with Pathwalking. J Struct Biol 196(3):289–298.

22. Baker MR, Rees I, Ludtke SJ, Chiu W, & Baker ML (2012) Constructing and validating initial Calpha models from subnanometer resolution density maps with pathwalking. Structure 20(3):450–463.

23. Cowtan K (2006) The Buccaneer software for automated model building. 1. Tracing protein chains. Acta Crystallogr D Biol Crystallogr 62(Pt 9):1002–1011.

24. Lindert S, et al. (2009) EM-fold: De novo folding of alpha-helical proteins guided by intermediate-resolution electron microscopy density maps. Structure 17(7):990–1003.

25. Wang RY, et al. (2015) De novo protein structure determination from near-atomic-resolution cryo-EM maps. Nat Methods 12(4):335–338.

26. Terashi G & Kihara D (2018) De novo main-chain modeling for EM maps using MAINMAST. Nature Communications 9(1).

27. Terwilliger TC, Adams PD, Afonine PV, & Sobolev OV (2018) A fully automatic method yielding initial models from high-resolution cryo-electron microscopy maps. Nat Methods 15(11):905–908.

28. Terashi G, Wang X, Prasad D, Nakamura T, & Kihara D (2024) DeepMainmast: integrated protocol of protein structure modeling for cryo-EM with deep learning and structure prediction. Nat Methods 21(1):122–131.

29. He J, Lin P, Chen J, Cao H, & Huang SY (2022) Model building of protein complexes from intermediate-resolution cryo-EM maps with deep learning-guided automatic assembly. Nat Commun 13(1):4066.

30. Zhang X, Zhang B, Freddolino PL, & Zhang Y (2022) CR-I-TASSER: assemble protein structures from cryo-EM density maps using deep convolutional neural networks. Nat Methods 19(2):195–204.

31. Pfab J, Phan NM, & Si D (2021) DeepTracer for fast de novo cryo-EM protein structure modeling and special studies on CoV-related complexes. Proc Natl Acad Sci U S A 118(2).

32. Giri N & Cheng J (2024) De novo atomic protein structure modeling for cryoEM density maps using 3D transformer and HMM. Nat Commun 15(1):5511.

33. Wang X, Zhu H, Terashi G, Taluja M, & Kihara D (2024) DiffModeler: large macromolecular structure modeling for cryo-EM maps using a diffusion model. Nat Methods.

34. Baek M, et al. (2021) Accurate prediction of protein structures and interactions using a three-track neural network. Science 373(6557):871–876.

35. Du Z, et al. (2021) The trRosetta server for fast and accurate protein structure prediction. Nature Protocols.

36. Ronneberger O, Fischer P, & Brox T (2015) U-Net: Convolutional Networks for Biomedical Image Segmentation. (Springer International Publishing), pp 234–241.

37. Su J, et al. (2024) RoFormer: Enhanced transformer with Rotary Position Embedding. Neurocomputing 568:127063.

38. Pintilie G, et al. (2020) Measurement of atom resolvability in cryo-EM maps with Q-scores. Nature Methods 17(3):328–334.

39. Davis IW, et al. (2007) MolProbity: all-atom contacts and structure validation for proteins and nucleic acids. Nucleic Acids Res 35(Web Server issue):W375–383.

40. Barad BA, et al. (2015) EMRinger: side chain–directed model and map validation for 3D cryo-electron microscopy. Nature Methods 12(10):943–946.

41. Abramson J, et al. (2024) Accurate structure prediction of biomolecular interactions with AlphaFold 3. Nature.

42. Turner J, et al. (2024) EMDB—the Electron Microscopy Data Bank. Nucleic Acids Research 52(D1):D456–D465.

43. Kui Xu Z-ED, Xing Zhang, Xin You, Pan Li, Nan Liu, Muzhi Dai, Chuangye Yan, Nieng Yan, Hong-Wei Wang, Sen-Fang Sui, Qiangfeng Cliff Zhang. (2024) Protein complex structure determination by structure prediction with cryo-EM density map constraints. https://github.com/kuixu/cryofold.

44. Zhang C, Shine M, Pyle AM, & Zhang Y (2022) US-align: universal structure alignments of proteins, nucleic acids, and macromolecular complexes. Nature Methods 19(9):1109–1115.

45. Amunts A & Nelson N (2009) Plant photosystem I design in the light of evolution. Structure 17(5):637–650.

46. Amunts A & Nelson N (2008) Functional organization of a plant Photosystem I: evolution of a highly efficient photochemical machine. Plant Physiol Biochem 46(3):228–237.

47. Amunts A, Ben-Shem A, & Nelson N (2005) Solving the structure of plant photosystem I--biochemistry is vital. Photochem Photobiol Sci 4(12):1011–1015.

48. Amunts A, Drory O, & Nelson N (2007) The structure of a plant photosystem I supercomplex at 3.4 A resolution. Nature 447(7140):58–63.

49. Wientjes E, Oostergetel GT, Jansson S, Boekema EJ, & Croce R (2009) The role of Lhca complexes in the supramolecular organization of higher plant photosystem I. J Biol Chem 284(12):7803–7810.

50. Lucinski R, Schmid VH, Jansson S, & Klimmek F (2006) Lhca5 interaction with plant photosystem I. FEBS Lett 580(27):6485–6488.

51. Storf S, Jansson S, & Schmid VH (2005) Pigment binding, fluorescence properties, and oligomerization behavior of Lhca5, a novel light-harvesting protein. J Biol Chem 280(7):5163–5168.

52. Storf S, Stauber EJ, Hippler M, & Schmid VH (2004) Proteomic analysis of the photosystem I light-harvesting antenna in tomato (Lycopersicon esculentum). Biochemistry 43(28):9214–9224.

53. Pan X, et al. (2018) Structure of the maize photosystem I supercomplex with light-harvesting complexes I and II. Science 360(6393):1109–1113.

54. Amunts A, Toporik H, Borovikova A, & Nelson N (2010) Structure determination and improved model of plant photosystem I. J Biol Chem 285(5):3478–3486.

55. Shekhar M, et al. (2021) CryoFold: Determining protein structures and data-guided ensembles from cryo-EM density maps. Matter 4(10):3195–3216.

56. Trabuco LG, Villa E, Schreiner E, Harrison CB, & Schulten K (2009) Molecular dynamics flexible fitting: A practical guide to combine cryo-electron microscopy and X-ray crystallography. Methods 49(2):174–180.

57. Lin TY, Goyal P, Girshick R, He K, & Dollar P (2020) Focal Loss for Dense Object Detection. IEEE transactions on pattern analysis and machine intelligence 42(2):318–327.

58. Li T, et al. (2024) All-atom RNA structure determination from cryo-EM maps. Nature Biotechnology.

59. Wang X, Terashi G, & Kihara D (2023) CryoREAD: de novo structure modeling for nucleic acids in cryo-EM maps using deep learning. Nature Methods 20(11):1739–1747.

60. Lin Z, et al. (2023) Evolutionary-scale prediction of atomic-level protein structure with a language model. 379(6637):1123–1130.

61. Shazeer N (2020) GLU Variants Improve Transformer. arXiv e-prints:2002.05202.

62. Glorot X, Bordes A, & Bengio Y (2011) Deep Sparse Rectifier Neural Networks. in Proceedings of the Fourteenth International Conference on Artificial Intelligence and Statistics, eds Geoffrey G, David D, & Miroslav D (PMLR, Proceedings of Machine Learning Research), pp 315--323.

## Supplementary References

1. Jamali K, et al. (2024) Automated model building and protein identification in cryo-EM maps. Nature 628(8007):450–457.

2. Kuhn HW (1955) The Hungarian method for the assignment problem. Naval Research Logistics Quarterly 2(1-2):83–97.

3. Ronneberger O, Fischer P, & Brox T (2015) U-Net: Convolutional Networks for Biomedical Image Segmentation. (Springer International Publishing), pp 234–241.

4. Sandler M, Howard A, Zhu M, Zhmoginov A, & Chen LC (2018) MobileNetV2: Inverted Residuals and Linear Bottlenecks. 2018 IEEE/CVF Conference on Computer Vision and Pattern Recognition, pp 4510–4520.

5. Gao SH, et al. (2021) Res2Net: A New Multi-Scale Backbone Architecture. IEEE Transactions on Pattern Analysis and Machine Intelligence 43(2):652–662.

6. Su H, et al. (2021) Improved Protein Structure Prediction Using a New Multi-Scale Network and Homologous Templates. Adv Sci (Weinh) 8(24):e2102592.

7. Lin Z, et al. (2023) Evolutionary-scale prediction of atomic-level protein structure with a language model. 379(6637):1123–1130.

8. Jumper J, et al. (2021) Highly accurate protein structure prediction with AlphaFold. Nature 596(7873):583–589.

9. Su J, et al. (2024) RoFormer: Enhanced transformer with Rotary Position Embedding. Neurocomputing 568:127063.

10. Burley SK, et al. (2023) RCSB Protein Data Bank (RCSB.org): delivery of experimentally-determined PDB structures alongside one million computed structure models of proteins from artificial intelligence/machine learning. Nucleic Acids Research 51(D1):D488–D508.

11. Terashi G, Wang X, Prasad D, Nakamura T, & Kihara D (2024) DeepMainmast: integrated protocol of protein structure modeling for cryo-EM with deep learning and structure prediction. Nat Methods 21(1):122–131.

12. Abramson J, et al. (2024) Accurate structure prediction of biomolecular interactions with AlphaFold 3. Nature.

13. Wú F, et al (2024) Structure of the II2-III2-IV2 mitochondrial supercomplex from the parasite Perkinsus marinus. bioRxiv:2024.2005.2025.595893

